# Molecular dynamics simulation of the truncated NMDA receptor in the open state

**DOI:** 10.1101/2025.09.15.676349

**Authors:** Xing Liu, Wenjun Zheng

## Abstract

The GluN1/GluN2A N-methyl-D-aspartate receptor (NMDAR) is a critical ligand-gated ion channel in the central nervous system, playing essential roles in synaptic plasticity, learning, and memory. Understanding its dynamics in the open/active state is paramount for deciphering its physiological functions and for developing targeted therapeutics. Despite many past efforts, the active/open state has not yet been fully resolved at atomic resolutions. To elucidate the molecular mechanism of the NMDAR activation, computer modeling and simulation are instrumental in providing detailed information about the dynamics and energetics of the receptor in various functional states. In this study, we started from a previously built open-state model of a truncated NMDAR, and then explored its energetics and dynamics with extensive molecular dynamics (MD) simulation (total simulation time is 5.6 µs). Based on the MD simulation, we employed an array of analysis tools to study the fluctuations/motions at the levels of individual residues, the channel pore, and the global structure, and identified a dynamic network of polar/nonpolar interactions between residues. Furthermore, we used machine learning to identify key interactions and residues relevant to channel opening/closing and validated them with evolutionary conservation grades and annotations of disease mutations in NMDAR. Taken together, our modeling and simulation provided rich structural and dynamic information which will guide functional studies of the activation of this key receptor.

## Introduction

The GluN1/GluN2A NMDA receptor is a heterotetrameric ion channel, typically composed of two GluN1 subunits and two GluN2A subunits, though triheteromeric assemblies also exist. It is activated by the co-agonists glycine (binding to GluN1) and glutamate (binding to GluN2A) [1–3] and the relief of magnesium block [4, 5]. Upon agonist binding, the receptor undergoes a series of conformational changes that lead to the opening of its central ion channel pore, allowing the influx of Ca2+ and Na+ ions, which are crucial for synaptic plasticity and excitatory neurotransmission [6]. The open/active state is characterized by a dilated pore that permits ion flow. This state is transient and is modulated by numerous allosteric modulators. Owning to its functional importance, various mutations in NMDA receptor are linked to neurological diseases, such as Alzheimer’s disease, depression, stroke, epilepsy and schizophrenia [7].

The architecture of NMDA receptors features three distinct layers of domains (Fig 1A-B): at the top are four amino-terminal domains (ATD) forming two heterodimers; below ATD are four ligand-binding domains (LBD) forming two heterodimers, below LBD is a tetrameric transmembrane domain (TMD). The two bi-lobed ATDs, each with R1 and R2 lobes, bind allosteric modulators, and regulate channel open probability and kinetics [8]. The two bi-lobed “clamshell” LBDs, each with S1 and S2 lobes, bind agonists and undergo closure of the S1-S2 cleft to trigger channel opening [9]. The TMD consists of four hydrophobic helical segments (M1 to M4) in each subunit, and the central M2/M3 helices line the channel pore [10, 11]. Within TMD, gating is mediated at least by two structural barriers. The primary gate (upper gate) is at the bundle crossing of the M3 helices (the classic “SYTANLAAF” motif region). Amin et al. [12] proposed a secondary gate (lower gate) at the apex of the M2 pore loops (i.e. N/Q/R site).

**Figure 1.**
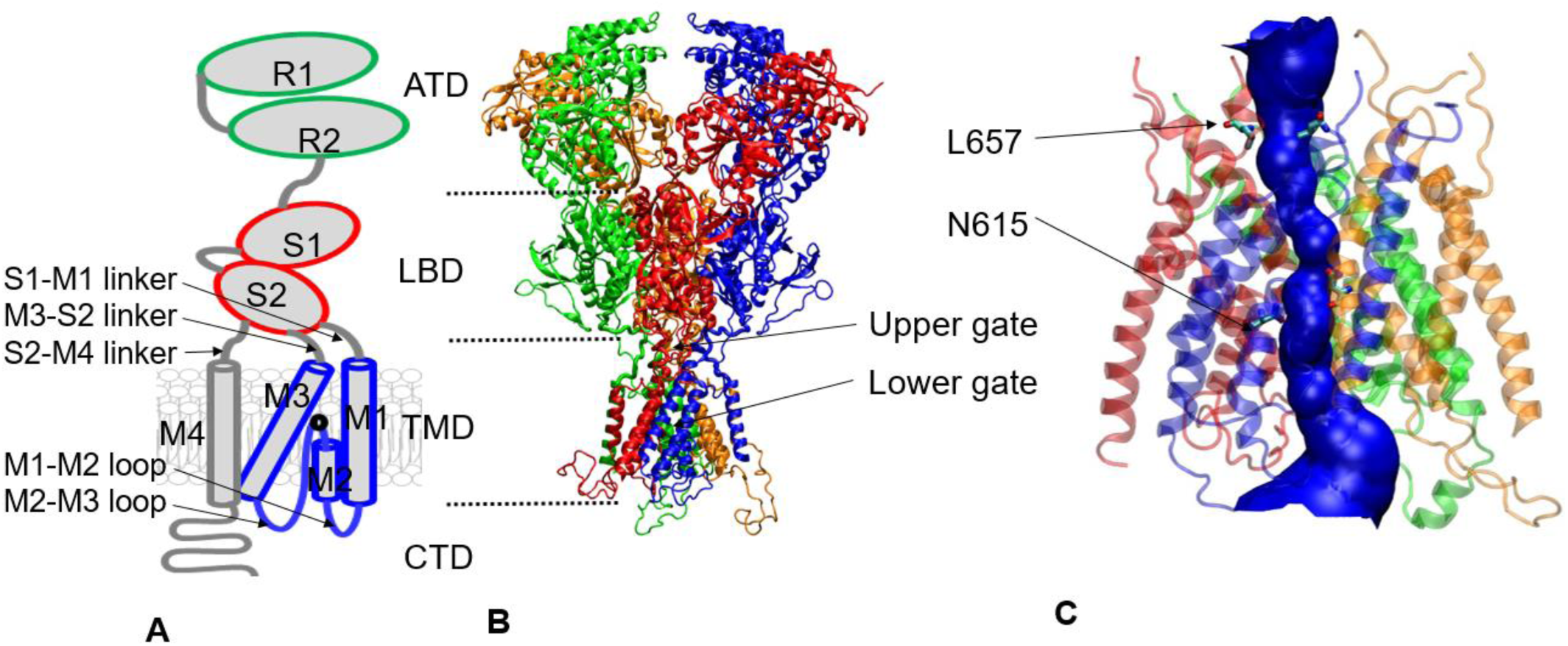
Structural view of NMDAR. (**A**) Subunit topology shows two external domains (ATD in green and LBD in red), each composed of two lobes (R1/R2 and S1/S2). Three LBD-TMD linkers (S1-M1, M3-S2, and S2-M4) connect the LBD with the TMD through M1, M3, and M4. (B) Structural model of a tetrameric N1/N2A receptor (without CTD), where the N2A subunits are colored in red/orange, and the N1 subunits are colored in blue/green, and the upper/lower gates are indicated. (C) From our MD simulation, a fully open channel pore is rendered as blue solid surfaces by the HOLE program: two representative residues, L657 at the upper gate and N615 at the lower gate, are shown.

Several structures of NMDA receptors have been solved by crystallography and cryo-electron microscopy (cryo-EM) at different resolutions. For example, two high-resolution crystal structures of heterotetrameric GluN1/GluN2B receptors are bound with agonists and allosteric inhibitors [13, 14], thus capturing an allosterically inhibited state [13–16]. Additionally, previous cryo-EM studies [17, 18] have visualized functional GluN1/GluN2B receptors at near/sub-nanometer resolutions in their active and antagonist-bound states, shedding lights on key conformational changes associated with activation [18] and antagonist inhibition [17]. More recently, Wang et al. reported a gallery of cryo-EM structures of the human GluN1-GluN2A NMDA receptor at an overall resolution of 4Å in complex with distinct ligands or modulators [19], showing that the binding of positive allosteric modulator shortens the distance between LBDs and TMD, which further stretches the opening of the gate. In another recent study, Chou et al. captured a GluN1–GluN2B receptor in the open state by cryo-EM [20] (with both agonists and a positive allosteric modulator bound). In a most recent cryo-EM study (Abbott et al. Cryo-EM snapshots of NMDA receptor activation illuminate sequential rearrangements, to be published in Science Advances), it was suggested that sequential bending of M3 helices opens the channel. Beyond GluN2A/B, cryo-EM studies have also captured other subtype structures of NMDAR. For example, Kang et al. solved the native tri-heteromeric GluN1–2B–2D receptor in the ligand-bound (activated) state, as well as inhibited and ketamine-blocked states [21]. Michalski et al. solved structures of GluN1–GluN3A bound to glycine versus antagonist with a 1-3-1-3 subunit arrangement [22].

Based on the structural studies, it was proposed that the transition to the open state involves substantial conformational rearrangements, including relative movement and rotation of the extracellular domains (ATDs and LBDs) with respect to the TMD. In particular, agonist binding drives the LBD to close its inter-lobe cleft, transmitting force via the GluN2 LBD–TMD linkers [20] to pull the pore-forming M3 helices outward, thus opening the channel gate. Notably, only when both glycine and glutamate bind do both LBDs fully close and exert enough tension to open the pore [20]. However, owning to the static nature of experimental structures and limited resolutions in certain key regions of NMDAR (e.g. in the TMD), the molecular details remain obscure on how conformational changes are transmitted from the extracellular ATD/LBD to channel pore upon receptor activation.

Computational methods have become indispensable for probing the dynamic transitions of the NMDA receptor, offering insights that complement experimental structural biology. These approaches range from atomistic MD simulations to coarse-grained models and kinetic analysis. MD is well-suited for simulating protein dynamics under physiological conditions at atomic resolution [23]. It nicely complements experimental dynamic measurements of NMDA receptor [24–28], and have provided atomistic insights into the gating mechanisms of NMDA receptor, elucidating conformational changes associated with the transition to the open/active state. Here are some highlights of previous MD simulation studies of NMDA receptors:

Dai et al. [29] used targeted MD simulations to develop an atomistic model for the open state of the GluN1/GluN2A NMDA receptor. They observed lobe closure of the ligand-binding domain produced outward pulling of the M3-D2 linkers, leading to outward movements of the C-termini of the pore-lining M3 helices and opening of the channel.

Amin et al. [12] showed that two separate gates mediate NMDA receptor activity and are under subunit-specific regulation: the lower gate (N-site) involving the M2 pore loop, and the upper gate at the M3 helical apex (bundle crossing). The two-gate model provides a kinetic explanation for the observation of bursts of openings/closings separated by long closures in single-channel-current measurements: GluN2-driven gating steps control entry/exit from bursts, whereas the GluN1 “inner gate” controls burst dwell times.

Černý et al. [30] conducted an aggregate of ∼1 µs of all-atom implicit membrane and solvent MD simulations of NMDAR from the open state to the closed state. They observed different gated states within the open state. They proposed that the ligand-induced rotation of extracellular domains is transferred by LBD–TMD linkers to the channel and responsible for its opening and closing.

Sinitskiy et al. [31] collected 30 μs of aggregated MD trajectories, observed rapid interconversions between 18 experimental atomic structures of NMDARs (i.e. multiple closed, open, and preopen states), and predicted 5 new conformations of the receptor.

Iacobucci et al. [32] performed MD simulation of NMDAR in the open state, and identified GluN1-I642 (on M3) and GluN2A-L550 (on the pre-M1 linker) form a new cross-subunit contact to stabilize the open state.

Despite progress in experimental and computational studies of NMDAR, several key questions about its open/active state remain unanswered. While the clamshell closure of LBD upon agonist binding and subsequent pulling of LBD-M3 linkers are well established, downstream rearrangements in TMD and channel gates are not clearly delineated. Although the "open state" was structurally characterized in two cryo-EM studies (see above), its structural heterogeneity as a dynamic ensemble remains to be fully explored. The nature of subunit-dependent channel gating, where GluN2A and GluN1 subunits play distinct roles, requires further investigations. The proposed secondary gate (lower gate) requires rigorous experimental/computational validations.

In this paper, we have employed an array of computational tools (including homology modeling, targeted and conventional MD simulation, channel pore analysis, principal component analysis, non-bonded energy calculations, and machine learning) to explore the structural dynamics of the NMDA receptor in the open state at the levels of individual residues, the channel pore, and the global tetrameric structure. Our goal is to map out a dynamic network of polar/nonpolar interactions between residues, and then identify key interactions and residues coupled with the channel pore opening/closing,

## Materials and methods

### Homology modeling

We have modeled a truncated NMDAR comprised of GluN1/GluN2A tetramer without ATD and CTD, containing the LBDs and TMD which are required for the essential channel function. For abbreviation, we refer to GluN1 as N1 and GluN2A as N2A in this study. Although the CTD is not included, its allosteric regulations of channel gating may be transmitted via M4 [33].

We downloaded a crystal structure of GluN1/GluN2B (PDB id: 4PE5) from Protein Data Bank (https://www.rcsb.org/). It captures an allosterically inhibited state of the receptor where the LBD clefts are closed (i.e. pre-activated) but the channel is closed. We obtained the amino acid sequences of *Rattus norvegicus* GluN1 and GluN2A (access ids: P35439 and Q00959) from the UniProt database [34] (https://www.uniprot.org/). We then submitted 4PE5 as template to the SWISS MODEL server [35] (https://swissmodel.expasy.org/) which generated a homology model for the above sequences. The resulting homology model includes chain A (residues 25-829 of GluN1), chain B (residues 29-839 of GluN2A), chain C (residues 25-832 of GluN1), and chain D (residues 31-836 of GluN2A). The missing residues were added by the SWISS MODEL server. We visualize protein structures using the VMD program (https://www.ks.uiuc.edu/Research/vmd/).

### MD simulation setup

Starting from a homology model of the GluN1/GluN2A receptor (see above), we truncated its ATDs to keep only LBDs and TMD. The truncated receptor includes GluN1 residues 397-838 and GluN2A residues 397-842. The purpose of truncation is to reduce the cost of MD simulation while retaining the core channel function.

To build the MD system, we submitted the truncated receptor structure to the Membrane Builder function [36–38] of the CHARMM-GUI webserver (https://www.charmm-gui.org/) which embeds the receptor in a bilayer of 1-palmitoyl-2-oleoyl phosphatidylcholine (POPC) lipids surrounded by a box of water, K^+^ and Cl^-^ ions (with 0.15 M concentration for the ions). The box contains 15-Å buffer of water/lipids extending from the receptor in each direction.

For MD system preprocessing, we used the NAMD program (https://www.ks.uiuc.edu/Research/namd/) to perform energy minimization and six steps of equilibration MD runs, which gradually reduce harmonic restraints applied to protein, lipids, water, and ions.

We ran targeted MD simulation to model the open state as described previously [32]. More specifically, we used an active-state conformation of LBD (PDB id: 5FXG, including GluN1 residues 397-441, 451-542, 667-798 and GluN2A residues 404-438, 453-537, 659-800) as the target of targeted MD, which was constructed from an active-state cryo-EM map (EMDB id: 3352). To drive conformational changes of the receptor to the active/open state (e.g. pulling the S2-M3 linkers of N2A open), we carried out a 30-ns targeted MD simulation using the Colvars module of NAMD (https://github.com/Colvars/colvars).

Following the above targeted MD simulation, we continued to run extensive conventional MD simulations. We restrained the LBD in its active-state conformation by applying 50 kj/mol/nm^2^ positional restraints to the Cα atoms. We used NAMD to conduct production MD runs in the NPT (isothermal-isobaric) ensemble with the Nosé-Hoover method [39, 40] at a temperature of 303 K, the Parrinello–Rahman method for pressure coupling [41], a 10-Å switching distance and a 12-Å cutoff distance for non-bonded interactions, and the particle mesh Ewald method for electrostatics calculations [42]. We constrained the hydrogen-containing bond lengths with the LINCS algorithm [43] to allow a 2-fs time step for the MD simulations. We used the CHARMM36 force field [44, 45] and TIP3P water model [46]. In total, we generated 7 independent MD trajectories of up to 800 ns each to sample the open-state conformational ensemble.

### RMSF analysis

To quantify the per-residue fluctuations of NMDAR during our MD simulation, we calculated the root mean square fluctuation (RMSF) as follows:

First, we saved 7x800 frames from 7 MD trajectories (i.e. one frame per ns) to build a conformational ensemble of the open state. Second, we superimposed the C_α_ coordinates of 640 residues in TMD and LBD-TMD linkers (residues 544-663 and 799-838 of N1, 539-658 and 803-842 of N2A) onto the initial structure with a minimal root mean square deviation (RMSD). Third, we calculated the following RMSF at residue position *n*: 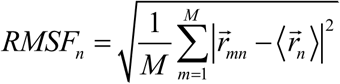, where 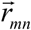 is the C_α_ position of residue *n* in frame 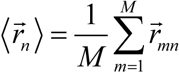 is the average C_α_ position of residue *n*, and *M=5600* is the total number of MD frames.

Finally, we averaged *RMSF_n_* for equivalent residues *n* of the N1/N2A dimer, and plotted RMSF as a function of residue position to show peaks and valleys corresponding to those regions with high and low fluctuations, respectively.

### Principal component analysis

To identify dominant modes of motions involved in the NMDAR channel dynamics, we performed the principal component analysis (PCA) as follows:

First, we superimposed the Cα coordinates of each MD frame onto the initial structure with a minimal RMSD. Second, we calculated a co-variance matrix comprised of the following 3×3 block matrices 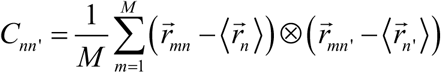 (see above for definitions of symbols). Third, we diagonalized the co-variance matrix and focused on top 10 PCA modes with the highest eigenvalues. Finally, we projected all MD frames along the eigenvectors of each PCA mode, and calculated their Pearson correlations with the minimal channel pore radius from the HOLE program (see below). We then visualized those few PCA modes correlated with channel pore open/closing (using the Normal Mode Wizard of VMD).

### Analysis of hydrogen bonds between residues

We find hydrogen bonds (HB) between any two polar non-hydrogen atoms (as acceptor and donor) with the donor-acceptor distance < 3.5 Å, and the donor-hydrogen-acceptor angle > 120°. We used the hydrogen bond plugin of the VMD program [47] to count pairwise HBs between residues in a MD frame (i.e. occu_ij_ is the number of HBs between residue i and j). To assess the relevance of those HBs to channel pore opening/closing, we calculated the Pearson correlation (PC) between occu_ij_ and the minimal channel pore radius (i.e. R_min_ from the HOLE program).

### Calculation of non-polar interaction energy between residues

We used the NAMD Energy plugin of VMD to calculate nonbonded interaction energy including the electrostatic energy and the van der Waals (vdW) energy between all pairs of residues (denoted vdW_ij_ for residues i and j). We focused on the vdW energy to describe nonpolar interactions between residues. We excluded those pairs with the maximal |vdW_ij_| < 1 kcal/mol. We calculated the Pearson correlation (PC) between vdW_ij_ and the minimal channel pore radius (i.e. R_min_ from the HOLE program). Then we aggregated PCs of pairwise vdW_ij_ to each residue i: PC_i_ = max(|PC| of vdW_ij_ for all j). We focused on residues with |PC_i_| > 0.3 and predicted them to be likely relevant to channel activation and involved in disease mutations [32]. Same aggregation of PCs was also performed for HB interactions.

We note the use of PC cutoff of 0.3 as rule of thumb for moderate-to-large correlations is common in the statistic field (see https://search.r-project.org/CRAN/refmans/effectsize/html/interpret_r.html). To further capture some nonlinear correlations, we have tried Spearman correlation instead of PC, but got very similar results.

### Residue conservation analysis

We submitted the sequences of GRIN2A (UniProt ID: Q12879) and GRIN1 (UniProt ID: Q05586) to the CONSURF server (http://consurf.tau.ac.il) described as follows:

Homologues Search: Homologues were collected from UNIREF90 database. Homologues search algorithm is HMMER. E-value cutoff is 0.0001. Number of Iterations is 1.

Homologues Thresholds: CD-HIT cutoff is 95% (This is the maximal sequence identity between homologues). Maximal number of final homologues is 200. Maximal overlap between homologues is 10% (If overlap between two homologues exceeds 10%, the highest scoring homologue is chosen). Coverage is 60% (This is the minimal percentage of the query sequence covered by the homologue). Minimal sequence identity with the query sequence is 35%.

Alignment, Phylogeny and Conservation Scores: Multiple Sequence Alignment was built using MAFFT. Phylogenetic tree was built using Neighbor Joining with ML distance. Conservation Scores were calculated with the Bayesian method. Amino acid substitution model was chosen by best fit.

Interpretation: Grade 8–9 indicate strongly conserved across homologs. This high conservation supports the idea the residue is functionally important.

### Pathogenic mutations of NMDAR residues

We obtained missense mutations/variations from the Clinvar database (https://www.ncbi.nlm.nih.gov/clinvar/) and disease mutations from the HGMD database (https://www.hgmd.cf.ac.uk/ac/all.php) for the human N1 gene (GRIN1) and N2A gene (GRIN2A). We focused on those mutations with disease relevance (e.g. with annotations ‘likely pathogenic’ or ‘pathogenic’ in Clinvar). For details, see the following files in Supplementary Information: hgmd_GRIN1_2025_pro, hgmd_GRIN2A_2025_pro, clinvar_result_grin1_2025, clinvar_result_grin2a_2025. We note that the sequences of the human N1 and N2A genes are identical to the corresponding rat genes in the truncated structure of TMD that we analyzed here.

### Machine learning to predict minimal pore radius using vdW/HB interactions

We conducted machine learning (ML) with RandomForestRegressor of scikit-learn. Our goal is to use pairwise vdw/HB interactions between residues as input features to predict the minimal channel pore radius (R_min_). We filtered out those weak vdW interactions with maximal value < 1 kcal/mol. We further removed those interactions with their ensemble standard deviation below a threshold value (denoted cutoff_std).

We split all 7x800 MD frames into a training set and a validation set. The former consists of first 600 frames of each trajectory, while the later includes the last 200 frames of each trajectory. We trained the regressor using the training set and performed hyperparameters tuning using the validation set.

We use the TreeExplainer from the shap python library (https://shap.readthedocs.io/) for calculating feature importance. SHAP (SHapley Additive exPlanations) in Python is a powerful library and framework for interpreting the predictions of machine learning models. It is based on Shapley values from cooperative game theory, adapted to explain individual predictions by attributing the contribution of each feature. We aggregated frame-wise shap values via summation, and then assigned importance to each residue i with the maximal aggregated shap value of those vdW/HB pairs involving residue i.

We used TPESampler of Optuna (https://optuna.org/) for hyperparameters optimization of RandomForestRegressor. 100 trials were taken for the following ranges of hyper-parameters: cutoff_std: [1.0, 0.5, 0.2, 0.1], max_depth: 3-10, max_features: [’sqrt’, ’log2’, 0.5, 0.2, 0.1], n_estimators: 100-1000.

## Results and discussion

### MD simulations of NMDAR in the open state

We previously used homology modeling and targeted MD simulation to build a model of NMDAR in a pre-activation state (with the LBD cleft closed and the M3-S2 linkers of N2A pulled outward) (for details, see Methods and [32]).

To further explore the conformational dynamics of NMDAR in the open state, we conducted 7 800-ns MD simulations at 30°C (with different initial random seeds), starting from a system of a truncated NMDAR model surrounded by a lipid bilayer and a box of water and ions (see Methods). While the NMDAR channel is known to be activated upon binding Glu and Gly in the LBD, possibly causing the LBD hinge bending to pull the M3-S2 linkers outward, it remains controversial how the TMD and the channel pore change structurally to enable rapid opening/closing. To address this issue, we combined the 7 MD trajectories to form a diverse ensemble of the open state for further analysis of its dynamics and energetics.

To evaluate the overall stability and dynamics of the MD simulation, we calculated the root mean squared deviation (RMSD) for all trajectories (relative to the initial model). The RMSD values initially increased and then stabilized in the range of 3-5Å (see Fig 2A), supporting the overall stability of the simulation despite the presence of highly flexible regions (e.g. the M1-M2 loop in the TMD). Nevertheless, the significant variations in RMSD between trajectories hint for a structurally heterogeneous open state of NMDAR which cannot be adequately described by few static snapshots and thus require extensive simulations and analysis at the ensemble scale.

**Figure 2.**
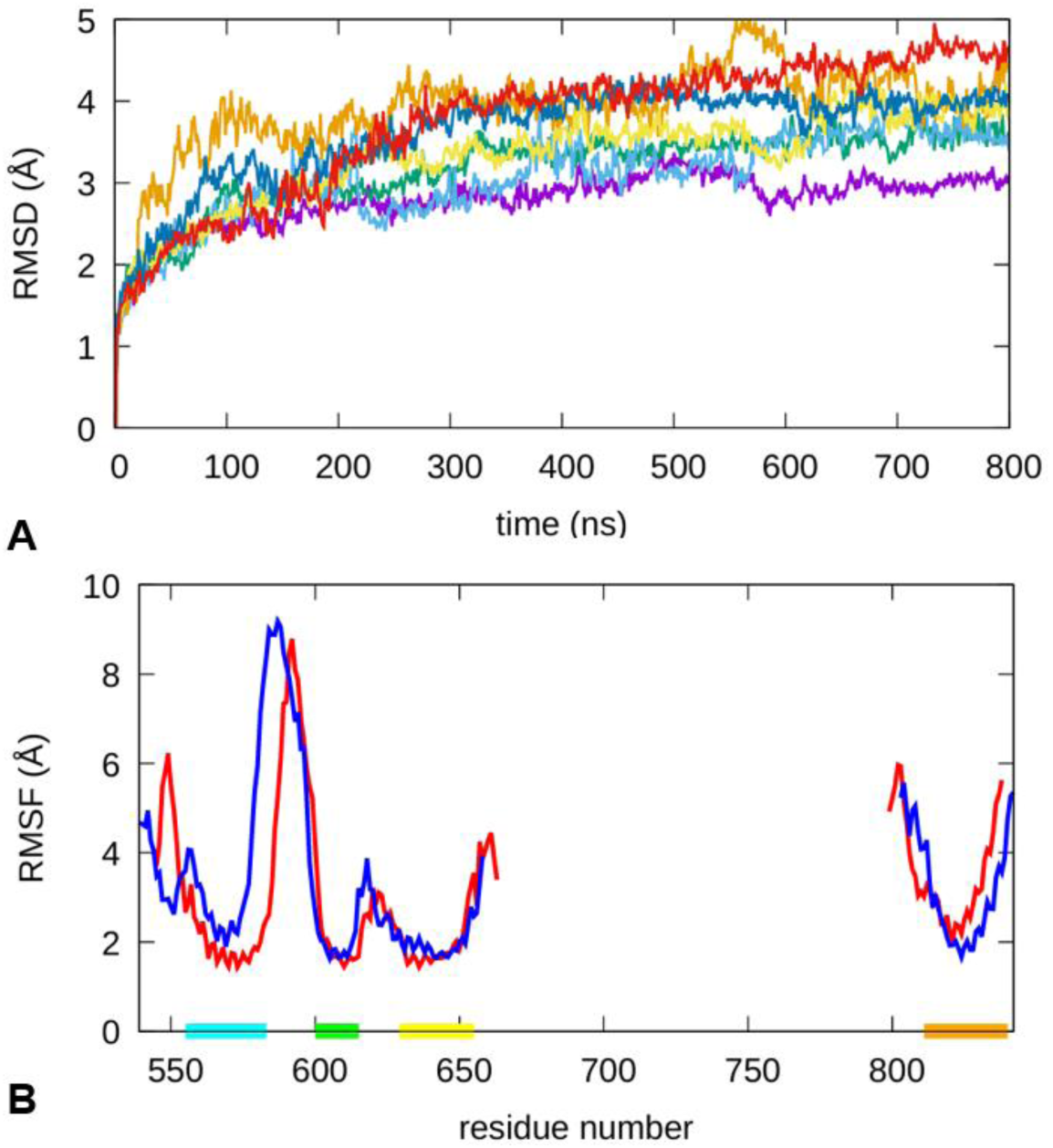
(A) RMSD of 7 MD trajectories of NMDAR. (B) RMSF for residues of N1 (red) and N2A (blue) based on the MD simulation. Ranges of residues corresponding to M1, M2, M3 and M4 are marked by horizontal bars colored in cyan, green, yellow, and orange, respectively. Residues of LBD S2 lobe are not shown because they are restrained in the MD simulation.

### RMSF analysis shows subunit-dependent flexibility in TMD and LBD-TMD linkers

To assess the conformational flexibility at the level of individual residues, we calculated the root mean square fluctuation (RMSF) in the open state ensemble (see Fig 2B). The RMSF profile exhibits pronounced peaks in the S1-M1 linker, M1-M2 loop, M2-M3 loop, M3-S2 linker, S2-M4 linker, and the C-terminus of the M4 helix (see Fig 2B). For example, the observed high flexibility of M1-M2 loop is consistent with its absence in experimentally solved structures of NMDAR. Previous studies of GluN2A_R586K variant [48] and phosphorylation [49] were mixed in ascertaining the functional roles of M1-M2 loop. In support of the functional importance of the above flexible loops/linkers, an early study found the conformational mobility of S1-M1 and S2-M4 linkers influence NMDAR gating [50].

To study how the N1/N2A asymmetry impacts their dynamics, we focused on the RMSF differences between the A/C (N1) and the B/D (N2A) subunits. While the RMSF of N1 and N2A agree well in the central TMD (i.e. M2 and M3), they differ in the periphery (in M1 and M4) and the various LBD-TMD linkers. Notably, N1 exhibits higher flexibility in the S1-M1 linker (near residue 550), the M3-S2 linker and the S2-M4 linker, while N2A shows higher flexibility in M1 and the M2-M3 loop. Such differences in dynamics may imply functional differences between N1 and N2A in regulating channel opening/closing. For example, the lower RMSF in the S1-M1 linker of N2A than N1 is consistent with a recent finding that the N1 residue I642 (on M3) and the N2A residue L550 (on the S1-M1 linker) form a cross-subunit contact to stabilize the open state [32].

Zooming into the lower and upper gate, the gate residues exhibit moderate fluctuations with RMSF between 2 and 3Å, and subtle differences between N1 and N2A: at the upper gate, A652 of N1 shows higher RMSF than the corresponding A650 of N2A (2.57 vs. 2.26 Å); but at the lower gate, N616 of N1 shows lower RMSF than the corresponding N614 of N2A (2.34 vs. 2.60 Å). These differences hint for functional differences between N1 and N2A subunits in controlling the two gates.

### HOLE analysis reveals a dynamic channel pore with two gates

To evaluate the degree of opening/closing of the NMDAR channel pore and quantify its dynamics in the MD simulations, we used the HOLE program [51] to calculate the pore radius at a series of points sampled along the pore axis for 5600 frames of the MD simulation. By inspecting the distribution of the minimal pore radius as a function of its nearest residue (see Fig 3A), we found two main clusters (circled in Fig 3A): one in M2 and M2-M3 loop, and the other in M3 and M3-S2 linker, corresponding to the lower gate and upper gate as proposed previously [12]. At the local maxima of R_min_ distribution, we identified residues N615 of N2A and L657 of N1 as two representative residues for the lower and upper gate, respectively, and assigned the minimum radius of those points nearest to these two residues as the radius of lower gate and upper gate, respectively (denoted R_lower_ and R_upper_).

**Figure 3.**
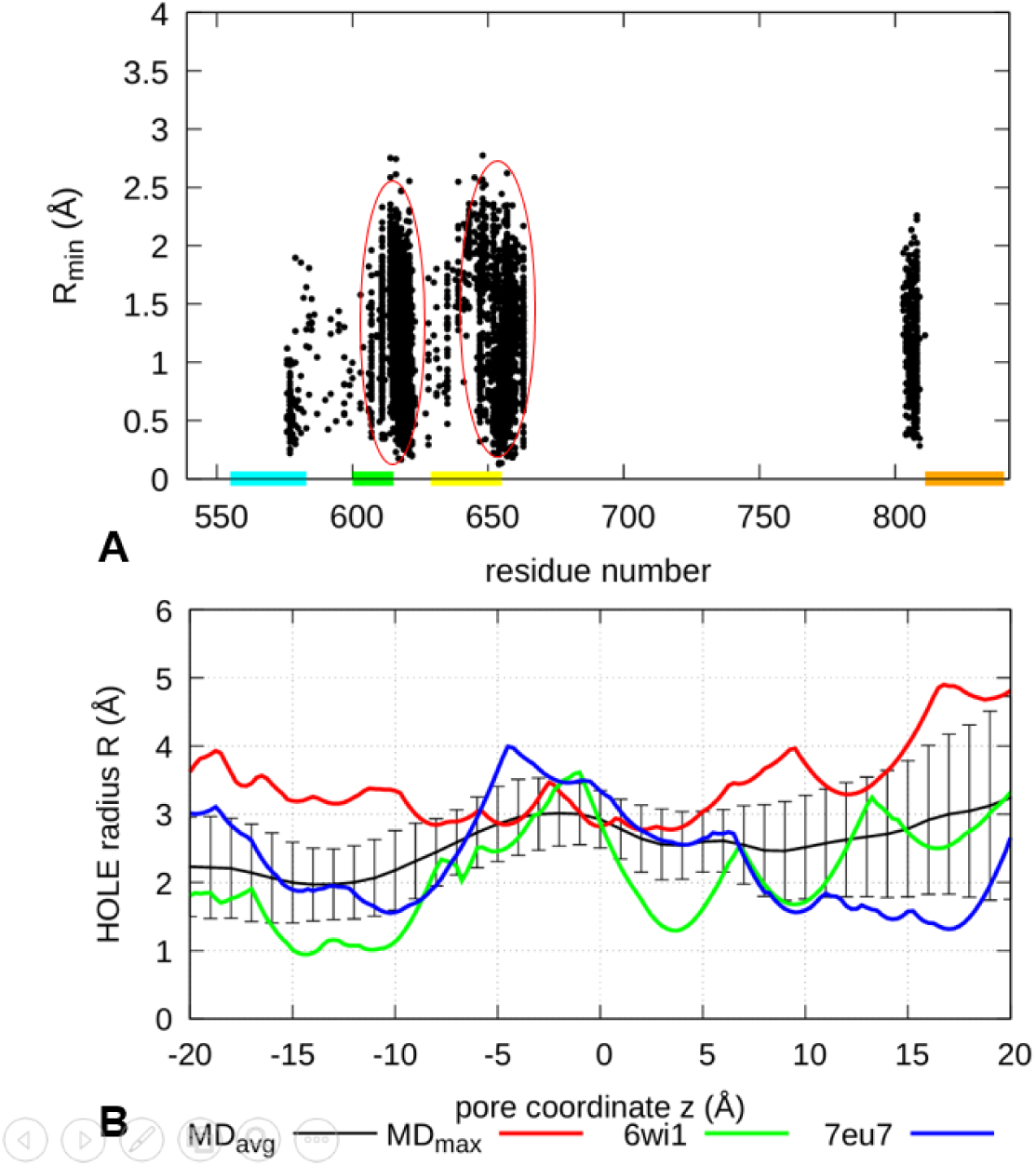
Results of HOLE analysis on MD frames. (**A**) Distribution of R_min_ over residues: for each frame, the HOLE program calculates the location with minimal radius (R_min_) along the channel pore and its nearest atom (identified by its residue number), corresponding to a data point in this plot. Two main clusters (corresponding to the lower and upper gates) are circled. Ranges of residues corresponding to M1, M2, M3 and M4 are marked by horizontal bars colored in cyan, green, yellow and orange, respectively. (**B)**. Channel pore radius profile: for each frame, the HOLE program calculates the pore radius R as a function of the z coordinate along the channel pore. Shown here are the pore radius profiles calculated for a representative frame with maximum R_min_ of 2.8Å (MD_max_), two experimental NMDAR structures (PDB ids: 6wi1 and 7eu7), and the ensemble average of pore radius profiles for all MD frames (MD_avg_) and the standard deviations as error bars.

We noted a wide distribution of R_min_ (0∼2.8Å) at both gates, suggesting a highly dynamic channel pore with rapid transient opening/closing of the channel in the open state. Based on the ensemble distribution of R_min_, we further estimated the probability of channel opening/closing by counting MD frames with R_min_ meeting the following criteria: a channel is deemed water permeable if *R*_min_ > 1.15 Å, likely/partially ion-conducting if

1.4 Å < *R*_min_ < 2.3 Å, and fully open if *R*_min_ > 2.3 Å [52]. Our MD simulation revealed a dynamic channel pore with 55% being water permeable, and 35% being partially open, and <1% being fully open (with maximum R_min_ of 2.8 Å).

We then visualized the pore radius profile along the channel axis. To clearly view the ensemble distribution of pore radius profiles, we showed the ensemble averages of pore radius (MD_avg_) and their standard deviations (see Fig 3B). The minimal radius of MD_avg_ (∼2Å) lies between z=-10 Å and z=-15 Å which is near the lower gate. For comparison, we calculated the pore radius profiles for an active-state structure (PDB id: 6wi1) and a ketamine-bound structure (PDB id: 7eu7). The former exhibits a more open upper channel (z>10 Å) while the latter shows a more open lower channel (z<-10 Å). In comparison, the MD_avg_ profile resembles 6wi1 for z>10 Å and 7eu7 for z<-10 Å, suggesting an overall more open pore than the experimental structures. We also showed the MD_max_ profile corresponding to a fully open channel (R_min_=2.8 Å) as sampled by the MD simulation (see Fig 1C).

Next, we tried to identify key factors (e.g., residues, interactions, motions, etc) that are coupled to the channel pore dynamics (as measured by R_min_).

### PCA reveals subunit-dependent structural changes correlated with channel opening/closing

To further extract dominant modes of structural fluctuations in the MD simulation, we applied the principal component analysis (PCA) to the MD-generated ensemble (see Methods), and focused on the top 10 PCA modes, which cumulatively account for 83% of total structural variations. The first PCA mode accounts for 22% of the total variations, while the remaining 9 modes each accounts for 1∼15%.

To identify those PCA modes correlated with the channel pore and the two gates, we calculated the Pearson correlations (PC) between the projections of MD frames along each mode and their R_min_, R_upper_, and R_lower_ (see Fig 4A). Among the top 10 PCA modes, mode 1 is highly correlated with R_lower_ (PC>0.4) and moderately correlated with R_upper_ (PC∼0.3); mode 4 is correlated with R_min_, R_upper_ and R_lower_ (PC>0.3); Mode 5 is highly correlated with R_min_ only.

**Figure 4.**
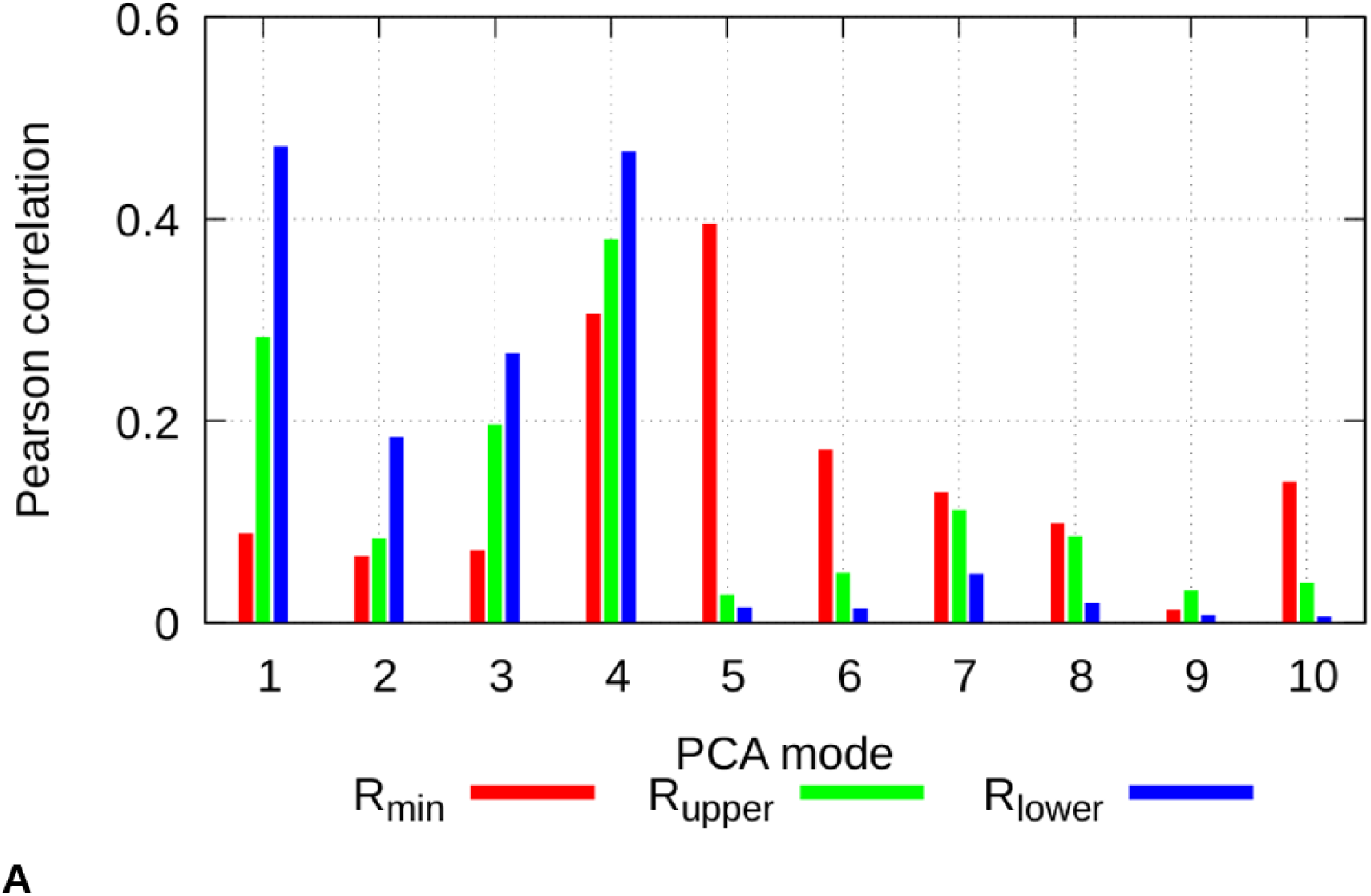
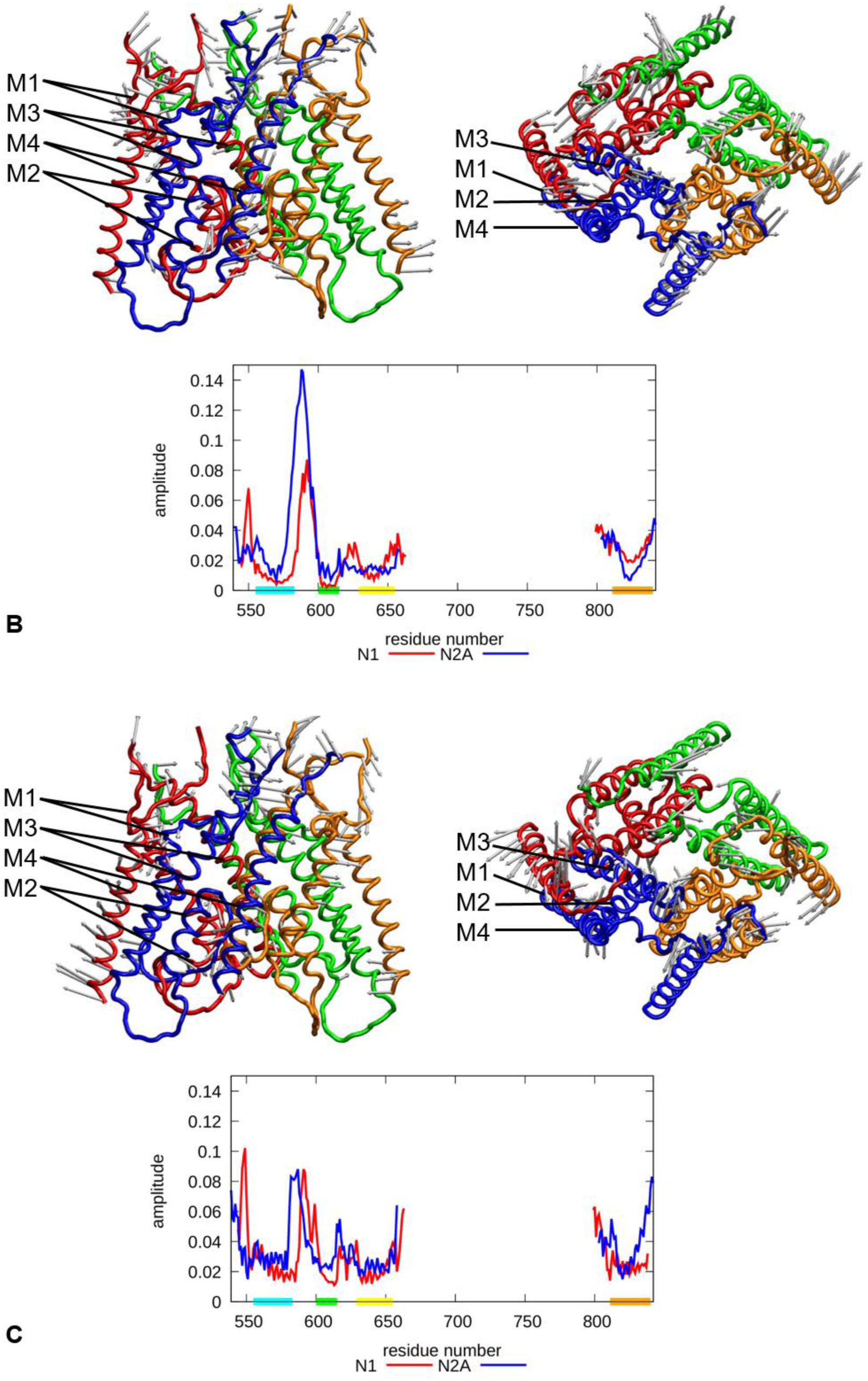
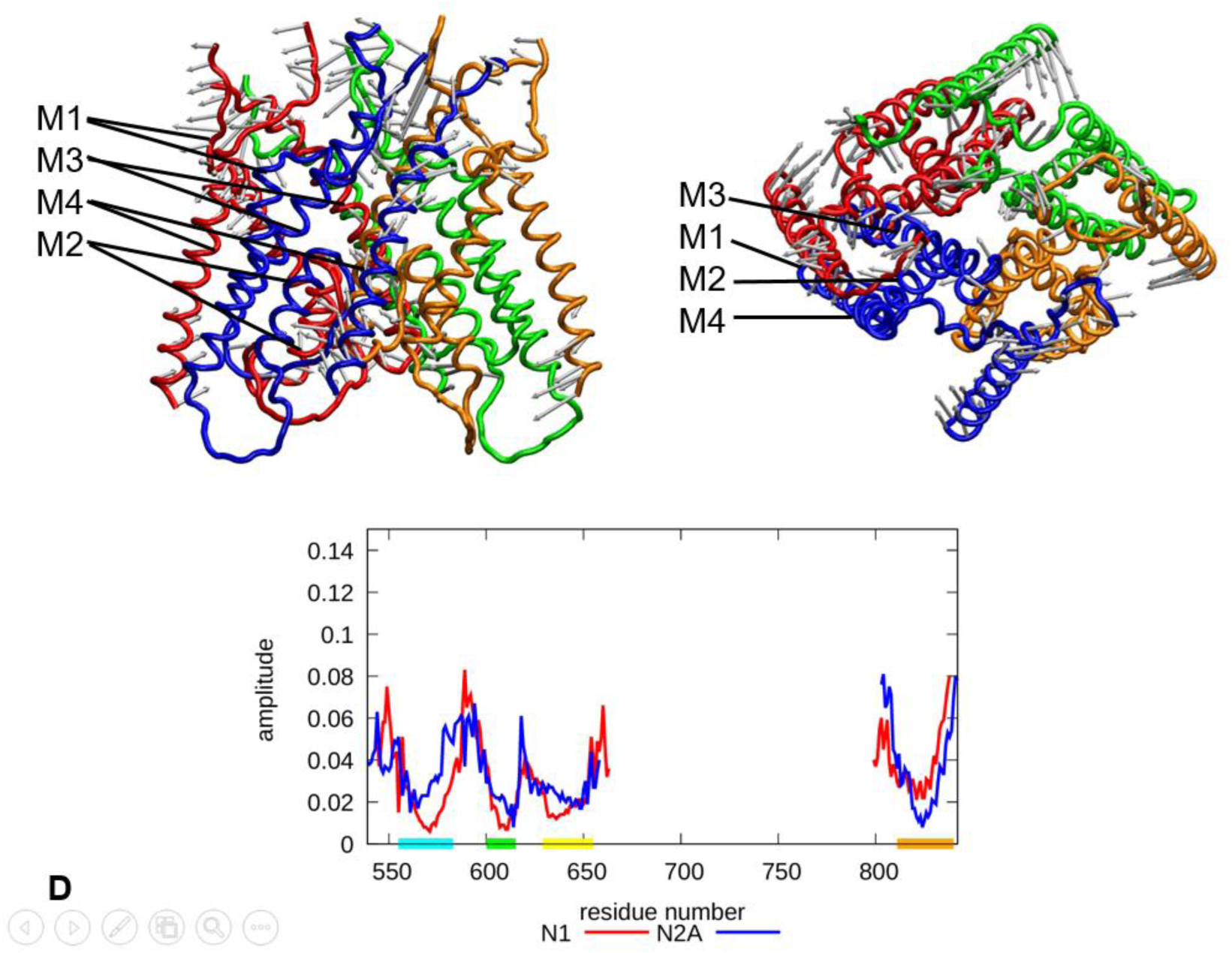
Analysis and visualization of PCA modes: (A). Assessing the relevance of top 10 PCA modes to channel gating: each MD frame is projected along a given PCA mode, and the Pearson correlations between these projections and R_min_/R_upper_/R_lower_ are shown for each mode. (B), (C) and (D) visualize the motions of PCA mode 1, 4, and 5 in side view (upper left panel) and top view (upper right panel), and plot its magnitude per residue (lower panel). Gray arrows indicate the motions of residues as described by the given mode.

The PCA mode 1 describes collective motions with subunit-dependent amplitude (Fig 4B): a spike in the S1-M1 loop of N1, prominent peaks in the M1-M2 loops of N1 and N2A, two minor peaks in the M2-M3 loop and the M3-S2 linker of N1, greater motions of M4 at the N/C-termini. Near the lower gate, N2A is more mobile than N1, hinting for a greater role of N2A in the gating of lower gate.

The PCA mode 4 also shows subunit-dependent amplitude of motions (Fig 4C): a spike in the S1-M1 loop of N1, major peaks in the M1-M2 loops of N1/N2A, a spike near the C-terminus of M2 in N2A, spikes in the M3-S2 linkers of N1/N2A, and pronounced peaks at the N-terminus of M4. At the lower gate, N2A is more mobile than N1, suggesting a greater role of N2A in the gating of lower gate. In contrast, near the upper gate, N2A and N1 exhibit similar amplitude of motions.

The PCA mode 5 also shows collective motions similar to the above 2 modes (Fig 4D), featuring greater motions in the S1-M1 loop, the M1-M2 loop, the M3-S2 linker, and the N/C-termini of M4. This mode could offer insight into how the NMDAR controls channel pore dynamics without involving the lower/upper gates.

### A network of polar/nonpolar interactions couple key residues to the channel pore

To find non-bonded polar interactions relevant to channel gating, we used the VMD program to identify all hydrogen bonds (HB) between pairs of residues in each MD frame (see Methods). We focused on those intra/inter-subunit HBs that are significantly correlated with the channel pore opening/closing (as measured by the Pearson correlations (PC) with R_min_) (see Fig 5 and Table 1). Those HB interactions with positive or negative correlations (i.e. PC>0.3 or PC<-0.3) are predicted to likely favor channel pore opening or closing.

**Figure 5.**
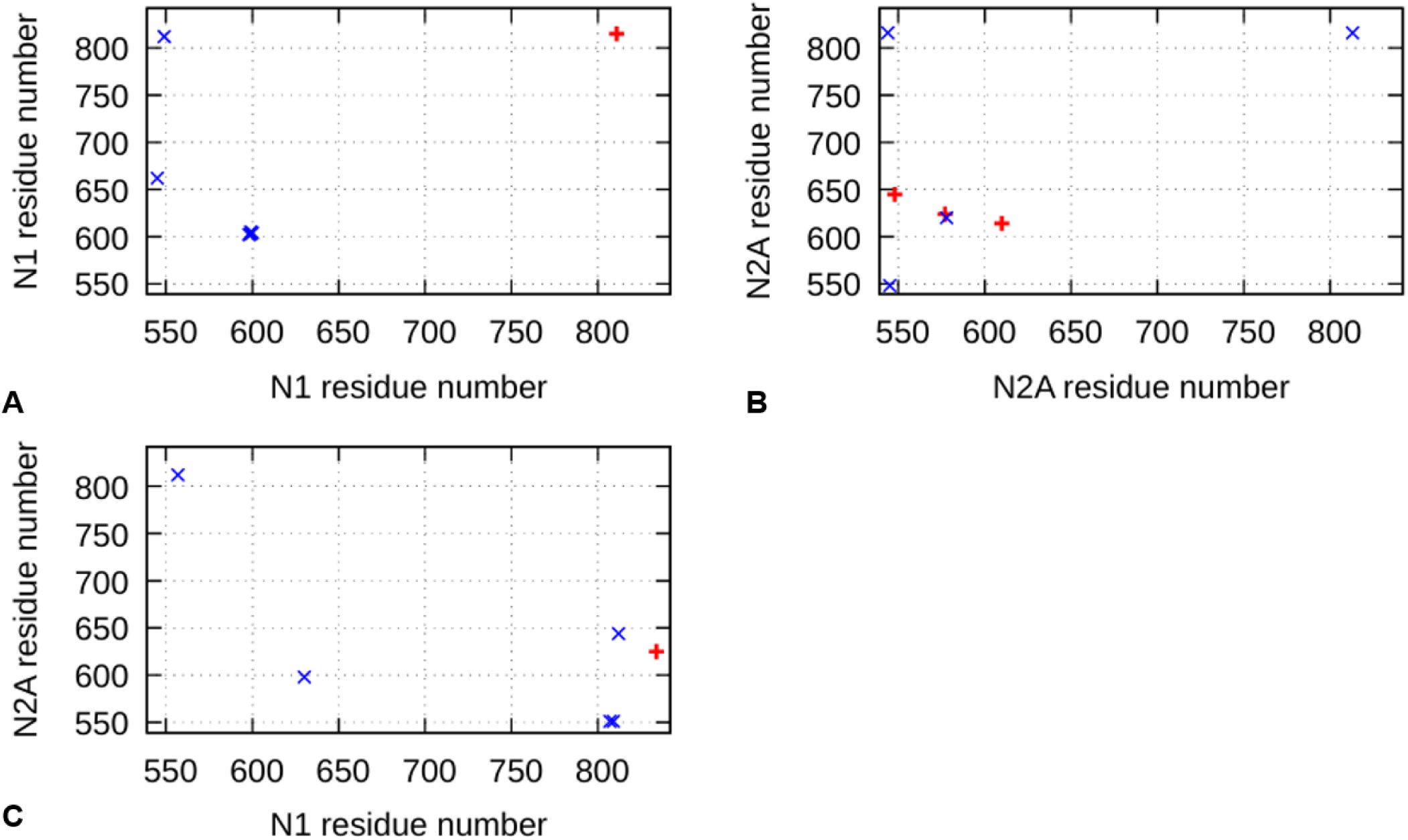
2D maps of intra/inter-subunit residue pairs that form HB interactions correlated with the minimal pore radius R_min_. Intra-N1, intra-N2A and N1-N2A residue pairs are shown in (A), (B) and (C). Residue pairs with positive PC>0.3 and negative PC<-0.3 are represented by red **+** and blue x, respectively.

**Table 1.** Residue pairs forming HB interactions correlated with the minimal pore radius R_min_ (with positive PC>0.3 or negative PC<-0.3).

| subunits involved | PC sign | residue pair | motifs involved |
| --- | --- | --- | --- |
| intra-N1 | positive | E811 G815 | between M4 and M4 |
|  | negative | E545 E662 | between S1-M1 linker and M3-S2 linker |
|  |  | S549 N812 | between S1-M1 linker and M4 |
|  |  | E598 T602 | between M1-M2 loop and M2 |
|  |  | D599 L603 | between M1-M2 loop and M2 |
|  |  | D599 S605 | between M1-M2 loop and M2 |
|  |  | A600 S604 | between M1-M2 loop and M2 |
| intra-N2A | positive | N614 G610 | between M2-M3 loop and M2 |
|  |  | G624 E577 | between M2-M3 loop and M1-M2 loop |
|  |  | Y645 A548 | between M3 and S1-M1 linker |
|  | negative | A548 S545 | between S1-M1 linker and S1-M1 linker |
|  |  | Q620 Y578 | between M2-M3 loop and M1-M2 loop |
|  |  | N816 V544<br>N816 D813 | between M4 and S1-M1 linker<br>between M4 and S2-M4 linker |
| N1-N2A | positive | E834 T625 | between M4 and M2-M3 loop |
|  | negative | P557 L812<br>R630 S598<br>T807 E551<br>T809 E551<br>N812 S644 | between S1-M1 linker and S2-M4 linker<br>between M2-M3 loop and M1-M2 loop<br>between S2-M4 linker and S1-M1 linker<br>between S2-M4 linker and S1-M1 linker<br>between M4 and M3 |

Only 5 HB-forming residue pairs were found to favor channel opening: Some of them are local interactions involved in the stability of local structures (such as E811-G815 and N614-G610, see Table 1). Other pairs are non-local interactions involved in global structural changes. For example, G624-E577 is between M2-M3 loop and M1-M2 loop in N2A, Y645-A548 is between M3 and S1-M1 linker in N2A, so they can affect the intra-N2A motions of M1/M2/M3. E834-T625 lies between M4 of N1 and M2-M3 loop of N2A, thereby controlling the inter-subunit motions that involve M2/M3/M4. Together, these HBs could help to stabilize specific intra/inter-subunit conformations that favor an open channel.

We found 15 HB-forming residue pairs that may favor channel closing (with PC<-0.3, see Fig5 and Table 1): Some of them are local interactions involved in the stability of local structures (such as A548-S545 in the S1-M1 linker of N2A, and N816-D813 in the M4 and S2-M4 linker of N2A). Other pairs form non-local interactions that may affect intra-subunit structural changes in N1 (e.g. E545-E662 and S549-N812) and in N2A (e.g. Q620-Y578 and N816-V544). 5 inter-subunit HB-forming pairs (i.e. P557-L812, R630-S598, T807-E551, T809-E551, N812-S644) may regulate motions between adjacent N1 and N2A subunits involving several key loops/linkers (see Table 1). Together, these intra/inter-subunit HB interactions could help to stabilize specific tertiary conformations that favor a closed channel.

In total, 35 key residues are involved in the above HBs (see Table 2). Here we discuss their functional significance. 33 of them are conserved (with Consurf grade of 8 or 9, see Methods): S549, P557, E598, D599, A600, T602, L603, S604, S605, R630, E662, T807, T809, N812, G815, E834 in N1; V544, S545, A548, E551, E577, Y578, S598, G610, N614, Q620, G624, T625, S644, Y645, L812, D813, N816 in N2A. This supports the functional importance of these residues. 11 of them are involved in disease mutations (as recorded in Clinvar and HGMD, see Methods): S549, P557, E662, T807, G815, E834 in N1; V544, A548, N614, S644, L812 in N2A.

**Table 2.** Key residues involved in HB interactions correlated with R_min_ Numerical mutation annotation as follows: 2 for pathogenic mutation in Clinvar or disease mutation in HGMD; 1 for mutation of uncertain significance or conflicting classifications in Clinvar; 0 for benign mutation in Clinvar.

| subunit | residue | mutation<br>annotation | Consurf<br>grade | motif |
| --- | --- | --- | --- | --- |
| N1 | E545 |  | 6 | S1-M1 linker |
|  | S549 | 2 | 9 | S1-M1 linker |
|  | P557 | 2 | 9 | S1-M1 linker |
|  | E598 |  | 8 | M1-M2 loop |
|  | D599 | 1 | 9 | M1-M2 loop |
|  | A600 |  | 9 | M1-M2 loop |
|  | T602 |  | 9 | M2 |
|  | L603 |  | 9 | M2 |
|  | S604 |  | 9 | M2 |
|  | S605 |  | 9 | M2 |
|  | R630 | 1 | 9 | M2-M3 loop |
|  | E662 | 2 | 9 | M3-S2 linker |
|  | T807 | 2 | 9 | S2-M4 linker |
|  | T809 |  | 9 | S2-M4 linker |
|  | E811 |  | 7 | M4 |
|  | N812 |  | 9 | M4 |
|  | G815 | 2 | 9 | M4 |
|  | E834 | 2 | 9 | M4 |
| N2A | V544 | 2 | 9 | S1-M1 linker |
|  | S545 |  | 9 | S1-M1 linker |
|  | A548 | 2 | 9 | S1-M1 linker |
|  | E551 | 1 | 8 | S1-M1 linker |
|  | E577 |  | 9 | M1-M2 loop |
|  | Y578 |  | 8 | M1-M2 loop |
|  | S598 |  | 8 | M1-M2 loop |
|  | G610 | 1 | 8 | M2 |
|  | N614 | 2 | 9 | M2-M3 loop |
|  | Q620 |  | 8 | M2-M3 loop |
|  | G624 |  | 9 | M2-M3 loop |
|  | T625 |  | 9 | M2-M3 loop |
|  | S644 | 2 | 9 | M3 |
|  | Y645 |  | 9 | M3 |
|  | L812 | 2 | 9 | S2-M4 linker |
|  | D813 |  | 9 | S2-M4 linker |
|  | N816 |  | 9 | M4 |

This suggests a possible mechanism for the pathogenesis of these mutations in disrupting the channel gating. Indeed, a functional study of disease-associated variants revealed that the S1–M1 linker of the NMDAR critically controls channel opening [53]. 23 of them are located in flexible loops/linkers (e.g. M1-M2 loop, M2-M3 loop, S1-M1 linker and S2-M4 linker), thus supporting the functional role of these flexible regions in channel gating.

To complement the above analysis of polar HB interactions, we further studied nonpolar interactions relevant to channel gating. To this end, we identified all van der waals (vdW) interactions between pairs of residues in each MD frame (see Methods). We focused on those intra/inter-subunit vdWs that are significantly correlated with the channel pore opening/closing (as measured by the Pearson correlations (PC) with R_min_) (see Fig 6 and Table 3). Those vdW interactions with positive or negative correlations (i.e. PC>0.3 or PC<-0.3) are predicted to likely favor channel pore opening or closing.

**Figure 6.**
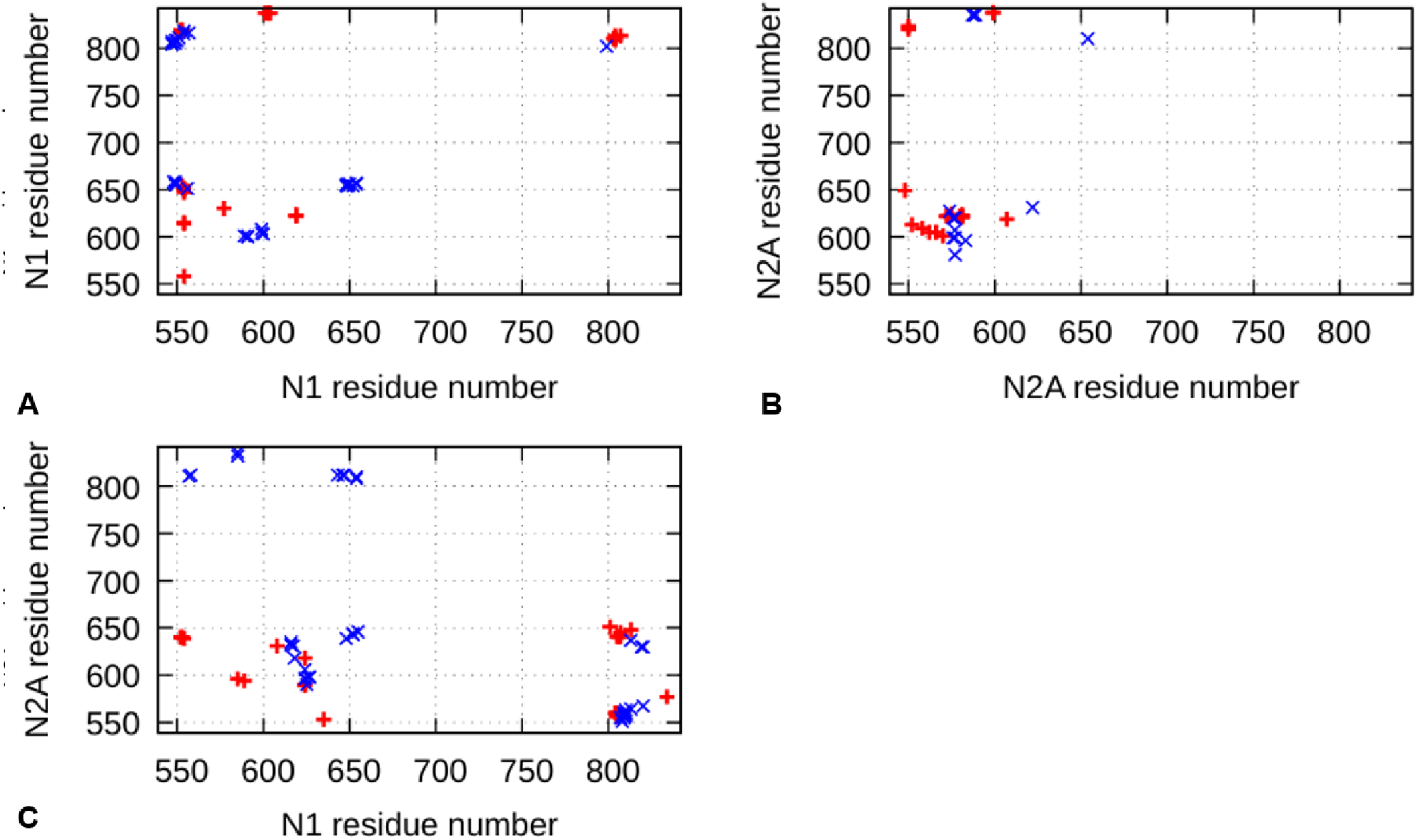
2D maps of intra/inter-subunit residue pairs that form vdW interactions correlated with the minimal pore radius R_min_. Intra-N1, intra-N2A and N1-N2A residue pairs are shown in (A), (B) and (C). Residue pairs with positive PC>0.3 and negative PC<-0.3 are represented by red **+** and blue x, respectively.

**Table 3.**
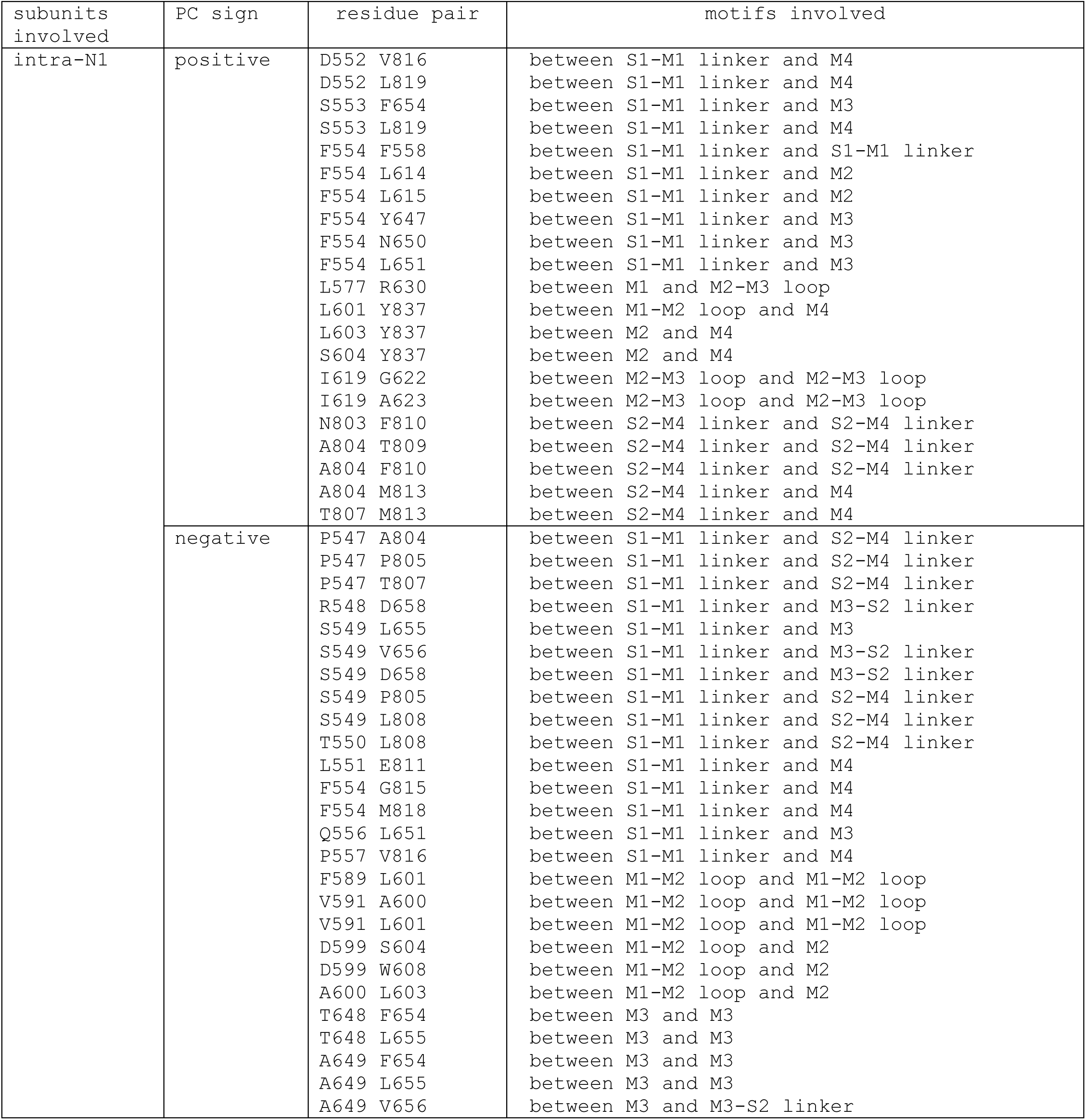
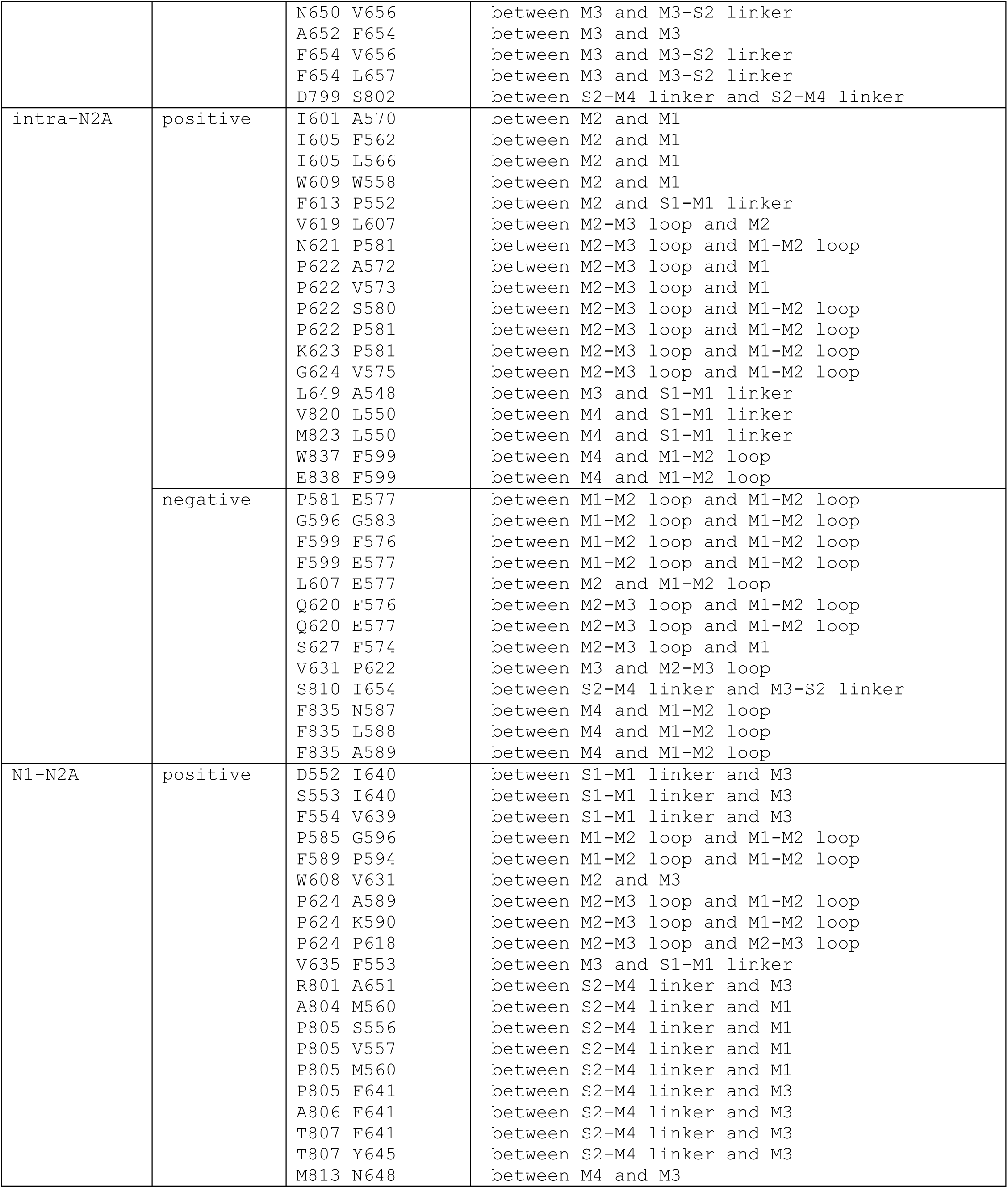
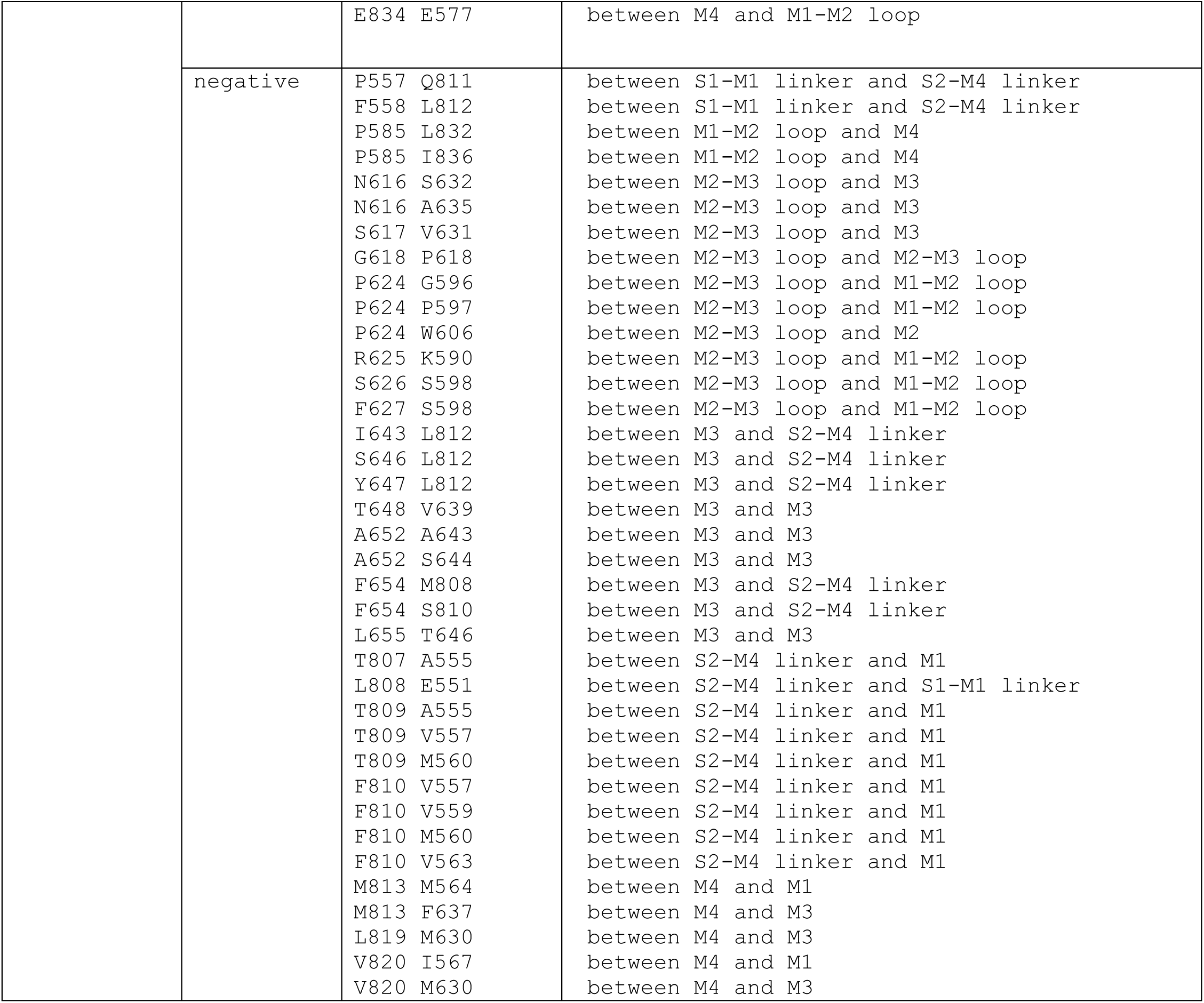
Residue pairs forming vdW interactions correlated with the minimal pore radius R_min_ (with positive PC>0.3 or negative PC<-0.3).

We first discussed those vdW-forming residue pairs that may interact to favor channel opening:

Among 21 intra-N1 residue pairs (see Table 3), 10 involve the S1-M1 linker and 5 involve the S2-M4 linker, supporting the importance of these LBD-TMD linkers to channel gating; 12 involve the pore-forming M2/M3 and associated loops, so they could directly affect channel dynamics; 19 involve the peripheral M1/M4 and associated loops, which may indirectly regulate channel dynamics via allostery.

Among 18 intra-N2A residue pairs (see Table 3), 8 involve the M2-M3 loop, and 7 involve the M1-M2 loop, enabling them to directly control the channel pore formed by M2/M3; 16 involve M2/M3 and associated loops which directly couple with channel dynamics; 17 involve M1/M4 and associated loops which allosterically regulate the channel pore.

Among 21 inter-subunit residue pairs (see Table 3), 9 involve the S2-M4 linker, 5 involve the M1-M2 loop, and 4 involve the S1-M1 linker, supporting a key role of these linkers/loops in inter-subunit couplings; 17 involve M2/M3 and associated loops, thus directly affecting channel dynamics; 19 involve M1/M4 and associated loops, hinting for their indirect allosteric couplings with the channel pore.

Next, we discussed those residue pairs that may interact to favor channel closing:

Among 31 intra-N1 residue pairs (see Table 3), 15 involve the S1-M1 linker, 7 involve the S2-M4 linker and 7 involve the M3-S2 linker, supporting a key role of these LBD-TMD linkers in channel gating; 20 involve M2/M3 and associated loops, while 22 involve M1/M4 and associated loops, suggesting the presence of both direct and indirect couplings to the channel pore.

Among 13 intra-N2A residue pairs (see Table 3), 7 involve the M3-S2 linker, 10 involve the M1-M2 loop, and 4 involve the M2-M3 loop, supporting the importance of these loops to channel gating; 13 involve M2/M3 and associated loops with direct couplings to the channel pore, while 12 involve M1/M4 and associated loops with allosteric couplings to the channel pore.

Among 37 inter-subunit residue pairs (see Table 3), 16 involve the S2-M4 linker, 10 involve the M2-M3 loop, and 7 involve the M1-M2 loop; 24 involve M2/M3 and associated loops, while 28 involve M1/M4 and associated loops.

By aggregating the above vdW interactions to individual residues (see Methods), we identified 144 key residues: 125 of them are conserved (with Consurf grade of 8 or 9, see Table 4), supporting their functional importance. 49 of them are involved in disease mutations (as recorded in Clinvar and HGMD, see Table 4) which may be explained by perturbation of channel dynamics. 75 of them are located in flexible loops/linkers (e.g. M1-M2 loop, M2-M3 loop, S1-M1 linker and S2-M4 linker), which supports the functional role of these flexible regions in mediating channel gating.

**Table 4.**
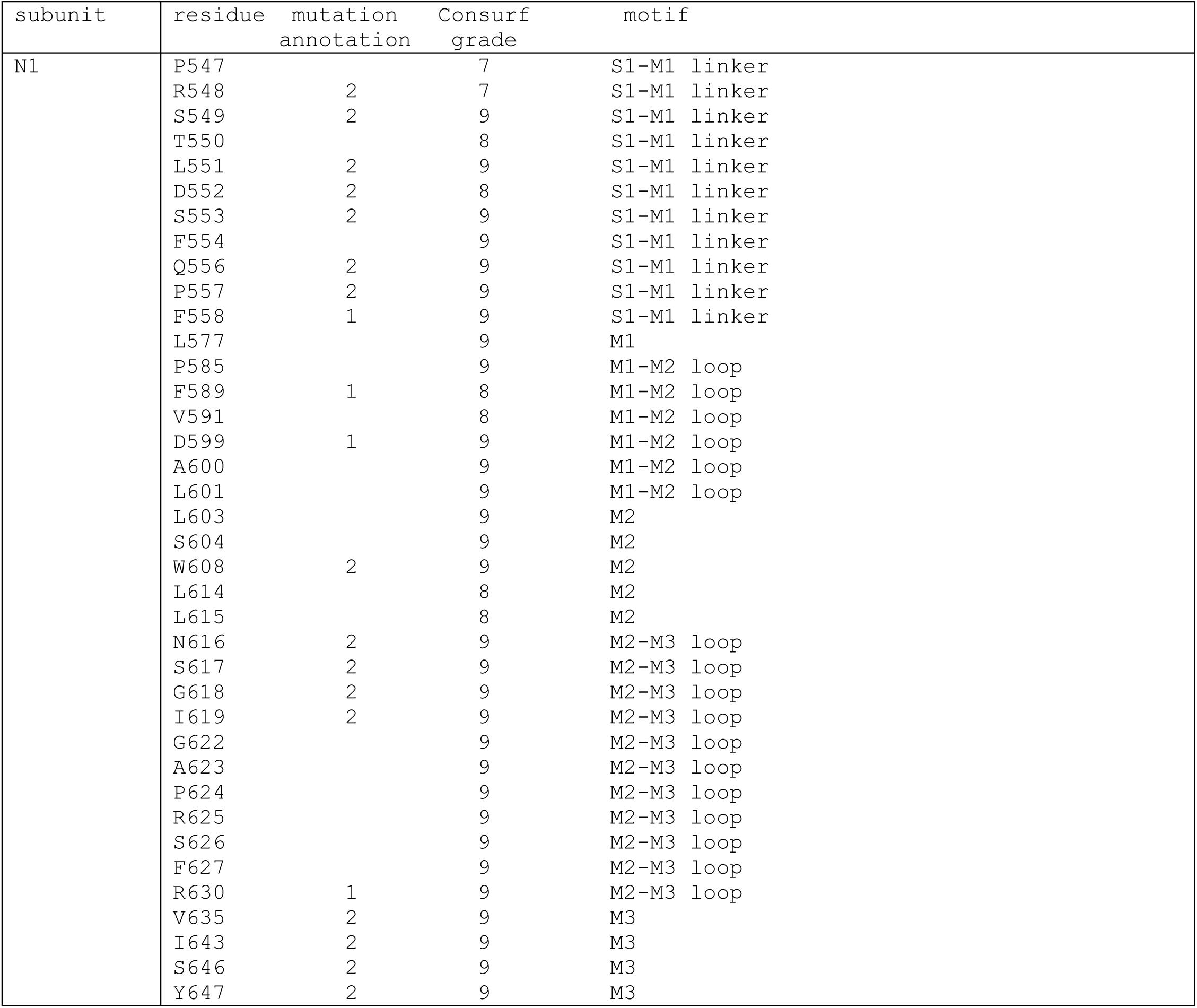
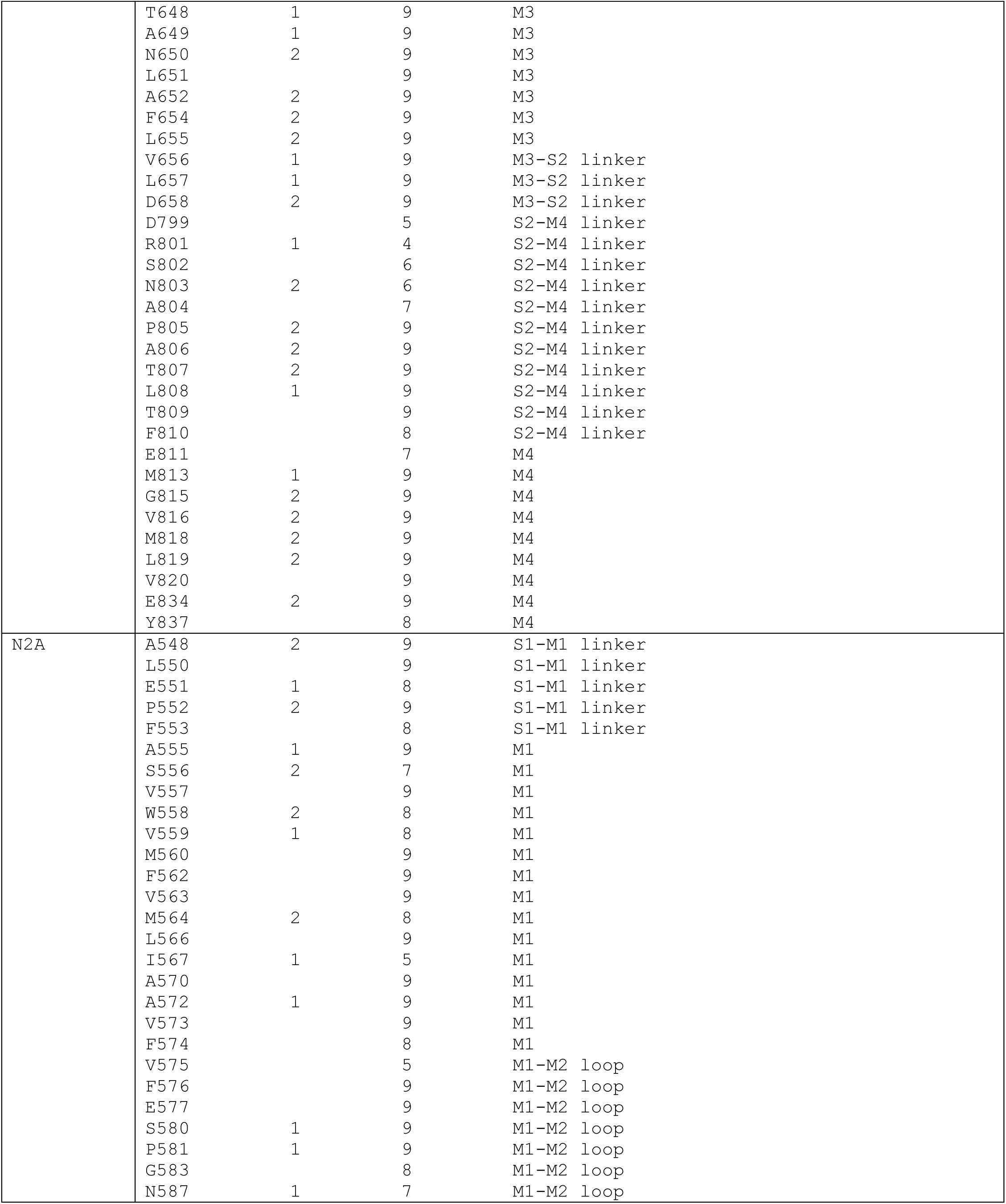
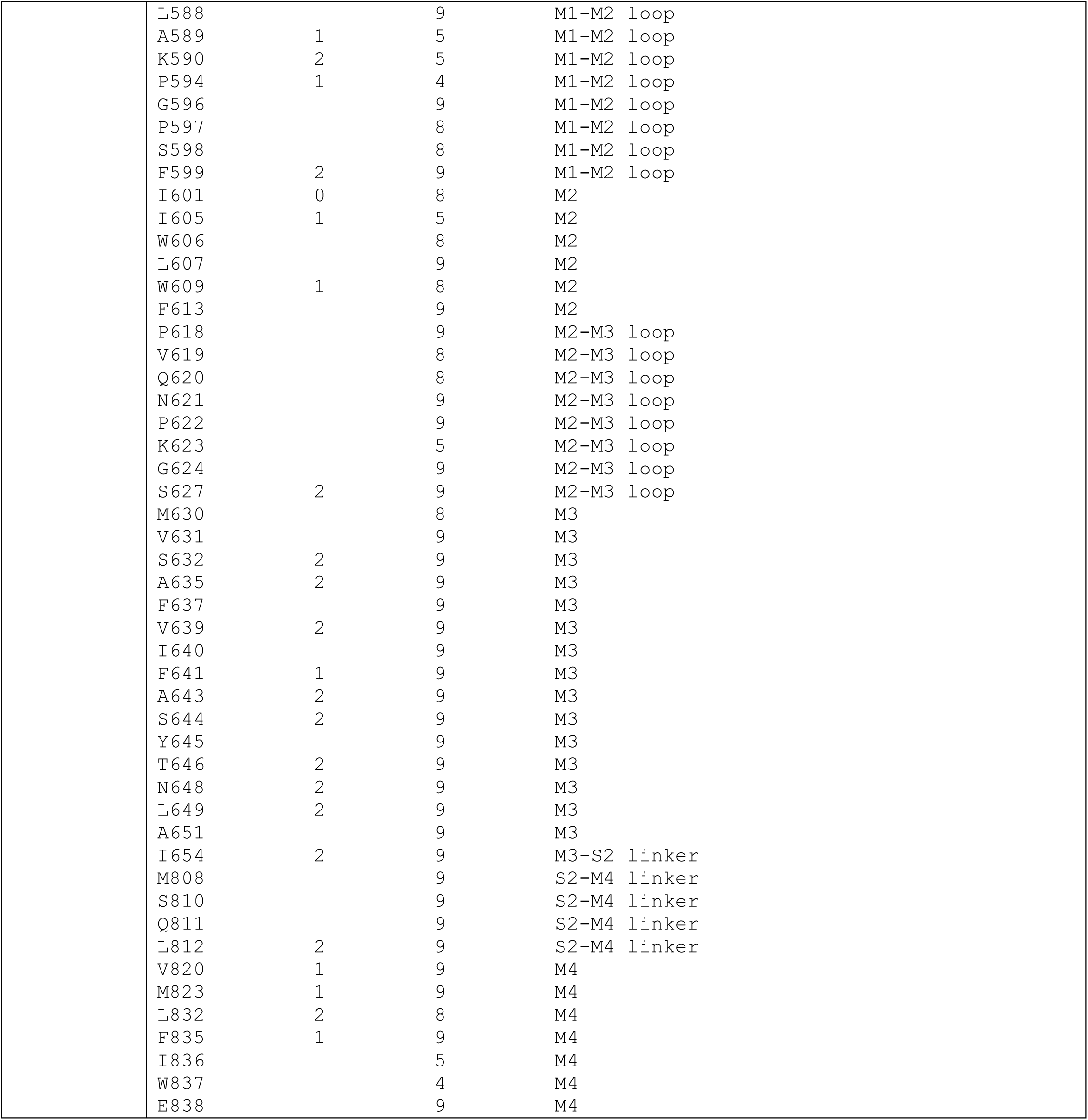
Key residues involved in vdW interactions correlated with R_min_. Numerical mutation annotation as follows: 2 for pathogenic mutation in Clinvar or disease mutation in HGMD; 1 for mutation of uncertain significance or conflicting classifications in Clinvar; 0 for benign mutation in Clinvar.

| subunit | residue | mutation<br>annotation | Consurf<br>grade | motif |
| --- | --- | --- | --- | --- |
| N1 | P547 |  | 7 | S1-M1 linker |
|  | R548 | 2 | 7 | S1-M1 linker |
|  | S549 | 2 | 9 | S1-M1 linker |
|  | T550 |  | 8 | S1-M1 linker |
|  | L551 | 2 | 9 | S1-M1 linker |
|  | D552 | 2 | 8 | S1-M1 linker |
|  | S553 | 2 | 9 | S1-M1 linker |
|  | F554 |  | 9 | S1-M1 linker |
|  | Q556 | 2 | 9 | S1-M1 linker |
|  | P557 | 2 | 9 | S1-M1 linker |
|  | F558 | 1 | 9 | S1-M1 linker |
|  | L577 |  | 9 | M1 |
|  | P585 |  | 9 | M1-M2 loop |
|  | F589 | 1 | 8 | M1-M2 loop |
|  | V591 |  | 8 | M1-M2 loop |
|  | D599 | 1 | 9 | M1-M2 loop |
|  | A600 |  | 9 | M1-M2 loop |
|  | L601 |  | 9 | M1-M2 loop |
|  | L603 |  | 9 | M2 |
|  | S604 |  | 9 | M2 |
|  | W608 | 2 | 9 | M2 |
|  | L614 |  | 8 | M2 |
|  | L615 |  | 8 | M2 |
|  | N616 | 2 | 9 | M2-M3 loop |
|  | S617 | 2 | 9 | M2-M3 loop |
|  | G618 | 2 | 9 | M2-M3 loop |
|  | I619 | 2 | 9 | M2-M3 loop |
|  | G622 |  | 9 | M2-M3 loop |
|  | A623 |  | 9 | M2-M3 loop |
|  | P624 |  | 9 | M2-M3 loop |
|  | R625 |  | 9 | M2-M3 loop |
|  | S626 |  | 9 | M2-M3 loop |
|  | F627 |  | 9 | M2-M3 loop |
|  | R630 | 1 | 9 | M2-M3 loop |
|  | V635 | 2 | 9 | M3 |
|  | I643 | 2 | 9 | M3 |
|  | S646 | 2 | 9 | M3 |
|  | Y647 | 2 | 9 | M3 |
|  | T648 | 1 | 9 | M3 |
|  | A649 | 1 | 9 | M3 |
|  | N650 | 2 | 9 | M3 |
|  | L651 |  | 9 | M3 |
|  | A652 | 2 | 9 | M3 |
|  | F654 | 2 | 9 | M3 |
|  | L655 | 2 | 9 | M3 |
|  | V656 | 1 | 9 | M3-S2 linker |
|  | L657 | 1 | 9 | M3-S2 linker |
|  | D658 | 2 | 9 | M3-S2 linker |
|  | D799 |  | 5 | S2-M4 linker |
|  | R801 | 1 | 4 | S2-M4 linker |
|  | S802 |  | 6 | S2-M4 linker |
|  | N803 | 2 | 6 | S2-M4 linker |
|  | A804 |  | 7 | S2-M4 linker |
|  | P805 | 2 | 9 | S2-M4 linker |
|  | A806 | 2 | 9 | S2-M4 linker |
|  | T807 | 2 | 9 | S2-M4 linker |
|  | L808 | 1 | 9 | S2-M4 linker |
|  | T809 |  | 9 | S2-M4 linker |
|  | F810 |  | 8 | S2-M4 linker |
|  | E811 |  | 7 | M4 |
|  | M813 | 1 | 9 | M4 |
|  | G815 | 2 | 9 | M4 |
|  | V816 | 2 | 9 | M4 |
|  | M818 | 2 | 9 | M4 |
|  | L819 | 2 | 9 | M4 |
|  | V820 |  | 9 | M4 |
|  | E834 | 2 | 9 | M4 |
|  | Y837 |  | 8 | M4 |
| N2A | A548 | 2 | 9 | S1-M1 linker |
|  | L550 |  | 9 | S1-M1 linker |
|  | E551 | 1 | 8 | S1-M1 linker |
|  | P552 | 2 | 9 | S1-M1 linker |
|  | F553 |  | 8 | S1-M1 linker |
|  | A555 | 1 | 9 | M1 |
|  | S556 | 2 | 7 | M1 |
|  | V557 |  | 9 | M1 |
|  | W558 | 2 | 8 | M1 |
|  | V559 | 1 | 8 | M1 |
|  | M560 |  | 9 | M1 |
|  | F562 |  | 9 | M1 |
|  | V563 |  | 9 | M1 |
|  | M564 | 2 | 8 | M1 |
|  | L566 |  | 9 | M1 |
|  | I567 | 1 | 5 | M1 |
|  | A570 |  | 9 | M1 |
|  | A572 | 1 | 9 | M1 |
|  | V573 |  | 9 | M1 |
|  | F574 |  | 8 | M1 |
|  | V575 |  | 5 | M1-M2 loop |
|  | F576 |  | 9 | M1-M2 loop |
|  | E577 |  | 9 | M1-M2 loop |
|  | S580 | 1 | 9 | M1-M2 loop |
|  | P581 | 1 | 9 | M1-M2 loop |
|  | G583 |  | 8 | M1-M2 loop |
|  | N587 | 1 | 7 | M1-M2 loop |

|  |  |  |  |
| --- | --- | --- | --- |
| L588 |  | 9 | M1-M2 loop |
| A589 | 1 | 5 | M1-M2 loop |
| K590 | 2 | 5 | M1-M2 loop |
| P594 | 1 | 4 | M1-M2 loop |
| G596 |  | 9 | M1-M2 loop |
| P597 |  | 8 | M1-M2 loop |
| S598 |  | 8 | M1-M2 loop |
| F599 | 2 | 9 | M1-M2 loop |
| I601 | 0 | 8 | M2 |
| I605 | 1 | 5 | M2 |
| W606 |  | 8 | M2 |
| L607 |  | 9 | M2 |
| W609 | 1 | 8 | M2 |
| F613 |  | 9 | M2 |
| P618 |  | 9 | M2-M3 loop |
| V619 |  | 8 | M2-M3 loop |
| Q620 |  | 8 | M2-M3 loop |
| N621 |  | 9 | M2-M3 loop |
| P622 |  | 9 | M2-M3 loop |
| K623 |  | 5 | M2-M3 loop |
| G624 |  | 9 | M2-M3 loop |
| S627 | 2 | 9 | M2-M3 loop |
| M630 |  | 8 | M3 |
| V631 |  | 9 | M3 |
| S632 | 2 | 9 | M3 |
| A635 | 2 | 9 | M3 |
| F637 |  | 9 | M3 |
| V639 | 2 | 9 | M3 |
| I640 |  | 9 | M3 |
| F641 | 1 | 9 | M3 |
| A643 | 2 | 9 | M3 |
| S644 | 2 | 9 | M3 |
| Y645 |  | 9 | M3 |
| T646 | 2 | 9 | M3 |
| N648 | 2 | 9 | M3 |
| L649 | 2 | 9 | M3 |
| A651 |  | 9 | M3 |
| I654 | 2 | 9 | M3-S2 linker |
| M808 |  | 9 | S2-M4 linker |
| S810 |  | 9 | S2-M4 linker |
| Q811 |  | 9 | S2-M4 linker |
| L812 | 2 | 9 | S2-M4 linker |
| V820 | 1 | 9 | M4 |
| M823 | 1 | 9 | M4 |
| L832 | 2 | 8 | M4 |
| F835 | 1 | 9 | M4 |
| I836 |  | 5 | M4 |
| W837 |  | 4 | M4 |
| E838 |  | 9 | M4 |

Together, the above residues (via their vdW interactions) form a network of direct and indirect couplings that control the dynamic balance between an open and a closed channel pore. These transiently forming/breaking vdW interactions (along with the HB interactions) drive global and local conformational changes that lead to rapid channel opening/closing in the open state.

Given that most TMD residues (273/320) are highly conserved, it is reasonable to expect many of them are relevant to the channel function. This may explain our findings of many residues involved in HB and vdW interactions significantly correlated with channel opening/closing. However, given the limited accuracy and time scales of our MD simulation, along with the simplicity/limitation of Pearson correlations, the above analysis cannot adequately capture complex nonlinear couplings to the channel dynamics. Additionally, the resulting large number of key residues/interactions as predicted above are not very informative in guiding experimental studies.

To address these caveats, we next employed machine learning algorithm to sharpen the predictions of a small number of residues most relevant to channel gating.

### Machine learning identifies important residues for channel gating

To identify key interactions and residues that contribute most to NMDAR channel opening/closing, we exploited machine learning (i.e. random forest) to predict R_min_ using pairwise vdW/HB interactions as input features, and then used an explainable AI method (i.e. shap) to assign feature importance (see Methods). Similar methods were published in previous studies [54]. Similar to the above analysis of Pearson correlations (PC) for HBs/vdWs, we aggregated the feature importance of pairwise interactions to individual residues (see Methods). Unlike the dense distributions of aggregated PCs for vdW/HB (see Fig 6A-D) where many residues show similarly high PCs, the distributions of aggregated feature importance are more sparse and spiky with few residues dominating (see Fig 7A-B). Therefore, the machine learning gave sharper predictions of key residues, allowing more focused follow-up studies.

**Figure 6.**
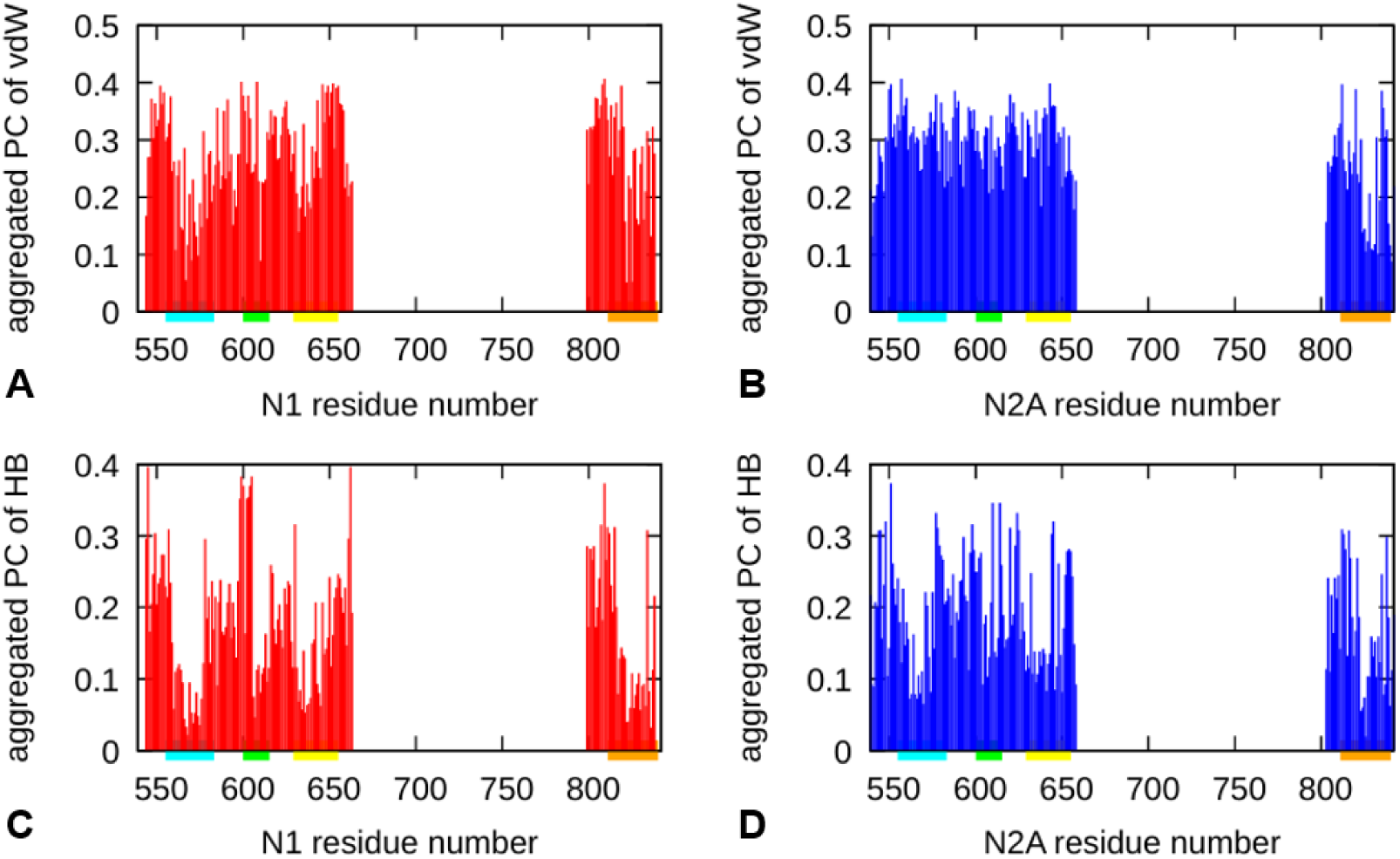
Aggregated Pearson correlations of pairwise vdW interactions (A-B), HB interactions (C-D) for residues of N1 and N2A.

**Figure 7.**
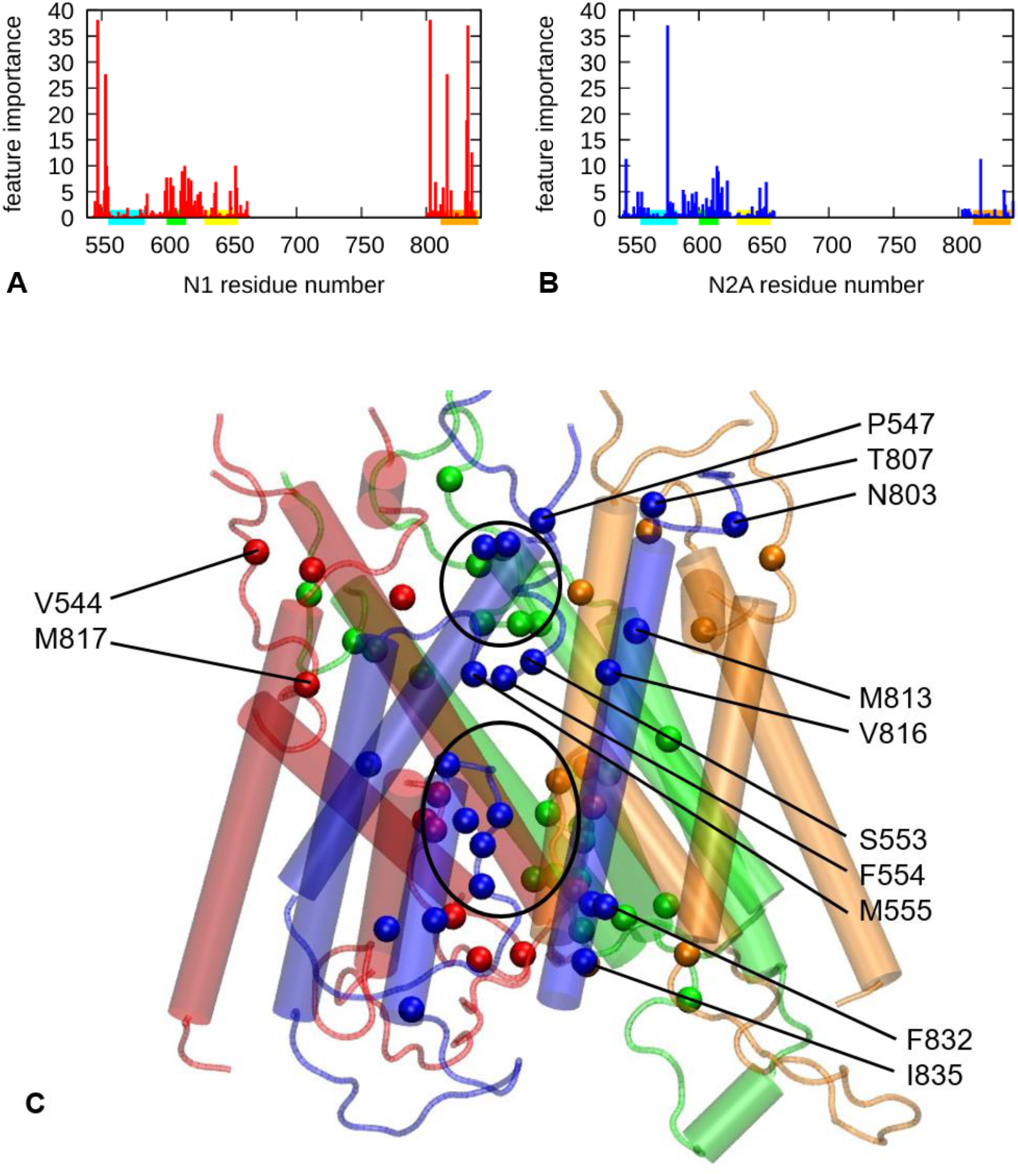
Aggregated feature importance from machine learning for residues of N1 and N2A is shown in (A-B). (C) shows predicted key residues (as ranked in top 10% by the aggregated feature importance) as balls colored by subunit (blue and green for N1, red and orange for N2A). The upper and lower gate regions are circled. Selected residues are labeled.

Next, we focused our discussion on the top 10% (32) residues as ranked by the aggregated feature importance. 29 of them are conserved (with Consurf grade of 8 or 9, see Table 5), supporting their functional importance; 19 of them are involved in disease mutations (as recorded in Clinvar and HGMD, see Table 5), hinting for a possible pathogenic mechanism associated with dysregulated channel gating; 16 of them are located in flexible loops/linkers (notably, M2-M3 loop, S1-M1 linker and S2-M4 linker), highlighting a key role of these dynamic regions in channel dynamics; 15 of them are on the peripheral M1/M4 and associated loops, so they may allosterically control channel gating (see Fig 7C).

**Table 5.** Predicted key residues as ranked top 10% by feature importance. Numerical mutation annotations are the same as Table 4

|  | residue | mutation<br>annotation | Consurf<br>grade | motif |
| --- | --- | --- | --- | --- |
| N1 | P547 |  | 7 | S1-M1 linker |
|  | S553 | 2 | 9 | S1-M1 linker |
|  | F554 |  | 9 | S1-M1 linker |
|  | M555 | 2 | 8 | S1-M1 linker |
|  | A600 |  | 9 | M1-M2 loop |
|  | L603 |  | 9 | M2 |
|  | S605 |  | 9 | M2 |
|  | G612 |  | 9 | M2 |
|  | L614 |  | 8 | M2 |
|  | S617 | 2 | 9 | M2-M3 loop |
|  | G618 | 2 | 9 | M2-M3 loop |
|  | I619 | 2 | 9 | M2-M3 loop |
|  | G638 | 2 | 9 | M3 |
|  | A653 | 2 | 9 | M3 |
|  | F654 | 2 | 9 | M3 |
|  | N803 | 2 | 6 | S2-M4 linker |
|  | T807 | 2 | 9 | S2-M4 linker |
|  | M813 | 1 | 9 | M4 |
|  | V816 | 2 | 9 | M4 |
|  | I831 |  | 9 | M4 |
|  | F832 |  | 6 | M4 |
|  | I835 | 2 | 8 | M4 |
| N2A | V544 | 2 | 9 | S1-M1 linker |
|  | F576 |  | 9 | M1-M2 loop |
|  | L611 | 2 | 9 | M2 |
|  | N614 | 2 | 9 | M2-M3 loop |
|  | N615 | 2 | 9 | M2-M3 loop |
|  | P618 |  | 9 | M2-M3 loop |
|  | P622 |  | 9 | M2-M3 loop |
|  | A647 | 2 | 9 | M3 |
|  | F652 | 2 | 9 | M3 |
|  | M817 | 2 | 9 | M4 |

In contrast with the predicted key residues by the Pearson correlation analysis (see Table 2 and 4), the fewer machine learning based predictions (see Table 5) are more concentrated with disease mutations. In support of the novelty of the machine learning predictions, 12/32 were not predicted by Pearson correlation analysis, supporting the power of machine learning in identifying complex nonlinear correlations with R_min_.

## Conclusions

The quest to understand the open/active state of the GluN1/GluN2A NMDA receptor has greatly benefited from the synergistic integration of computational and experimental approaches. Cryo-electron microscopy has provided crucial high-resolution structural snapshots of the receptor in various liganded and modulated states. Kinetic analyses of single-channel activity have identified functional classes of conformers populated during activation [55]. Molecular dynamics simulations hold promise of bridging the gap between structures and function, elucidating the dynamic fluctuations of residues, predicting intermediate conformations, and identifying critical intra/inter-subunit interactions at an atomistic level. In this study, our MD simulations of the open state have made progress toward shedding new lights on the dynamics of key parts of the NMDAR, including various LBD-TMD linkers, transmembrane helices and associate loops. Our simulation has revealed a highly dynamic channel pore undergoing rapid opening/closing at two gates, coupled with a dynamic network of polar/nonpolar interactions. Furthermore, we have used machine learning to predict important residues and validate them with high conservation and involvement in disease mutations, which will make promising targets for future experimental studies of channel gating of NMDAR.

## Acknowledgements

We thank Prof. Popescu for reading and commenting on the manuscript. We also thank UB CCR for providing computing resources. The MD trajectories, interaction data, and machine learning script will be shared on Zenodo (https://zenodo.org/uploads/17096455)

